# Histological aging signatures enable tissue-specific disease prediction from blood

**DOI:** 10.1101/2024.11.14.618081

**Authors:** Ernesto Abila, Iva Buljan, Yimin Zheng, Tamas Veres, Zhilong Weng, Maja Nackenhorst, Wolfgang Hulla, Yuri Tolkach, Adelheid Wöhrer, André F. Rendeiro

**Affiliations:** CeMM Research Center for Molecular Medicine of the Austrian Academy of Sciences, Lazarettgasse 14 AKH BT 25.3, 1090, Vienna, Austria; Ludwig Boltzmann Institute for Network Medicine at the University of Vienna, Augasse 2-6, A-1090 Vienna, Austria; Institute of Pathology, University Hospital Cologne, Cologne, Germany; Department of Pathology, Medical University of Vienna, Vienna, Austria; Institute of Pathology, Landesklinikum Wiener Neustadt, Wiener Neustadt, Austria; Division of Neuropathology and Neurochemistry, Department of Neurology, Medical University of Vienna, Vienna, Austria; Department of Pathology, Neuropathology and Molecular Pathology, Medical University of Innsbruck, Innsbruck, Austria

**Keywords:** aging, histopathology, deep learning, aging clocks, tissue biology, biomarkers

## Abstract

Aging, the leading risk factor for numerous diseases, manifests through diverse structural and architectural changes in human tissues, providing an opportunity to quantify and interpret tissue-specific aging. To address this, we present a comprehensive assessment of tissue changes occurring during human aging, utilizing a vast array of whole slide histopathological images from the Genotype-Tissue Expression Project (GTEx), primarily reflecting non-diseased tissue samples. Using deep learning, we analyzed 25,712 images from 40 distinct tissue types across 983 individuals, quantifying nuanced morphological changes that tissues undergo with age.

We developed ‘tissue clocks’—predictors of biological age based on tissue images—which achieved a mean prediction error of 4.9 years. These clocks were associated with established aging markers, including telomere attrition, subclinical pathologies, and comorbidities. In a systematic assessment of biological age rates across organs, we identified pervasive non-uniform rates of aging across the human lifespan, with some organs exhibiting earlier changes (20-40 years old) and others showing bimodal patterns of age-related changes. We also uncovered several associations between demographic, lifestyle, and medical history factors and tissue-specific acceleration or deceleration of biological age, highlighting potential modifiable risk factors that influenced the aging process at the tissue level. Finally, by combining paired histological images and gene expression data, we developed a strategy to predict tissue-specific age gaps from blood samples. This approach was validated in independent cohorts covering eight diseases, ranging from acute conditions like stroke to chronic diseases such as cystic fibrosis and Alzheimer’s disease. It successfully recovered significant associations with disease-relevant organs and revealed patterns of systemic and tissue-specific aging that may reflect broader physiological changes in health and disease.

This work offers a new perspective on the aging process by positioning tissue structure as an integrator of cellular and molecular changes that reflect the physiological state of organs in health and disease. It underscores the value of histopathological imaging as a tool for understanding human aging and provides a foundation for the monitoring of tissue-specific aging processes in age-associated diseases.

## Introduction

Aging is a fundamental biological process with far-reaching implications for human health and disease^1^. Despite its inevitability, the mechanisms underlying aging and the resulting age-related changes display striking diversity across tissues^2,3^. This heterogeneity presents a considerable challenge in elucidating the pathways through which aging influences tissue function decline and disease onset. Contributions from evolutionary biology^4–6^, cellular biology and molecular genetics^7–9^ have enriched our understanding of the genetic and environmental factors that influence the human body throughout the lifespan. However, translating these insights into a coherent understanding of the aging process, which is associated with changes in physiological function at the organism level, remains a considerable challenge^10–12^.

Research into human aging has historically faced several limitations. Animal models, despite their value, fall short of capturing the complexity of human aging due to physiological and lifespan differences, as well as divergent environmental exposures^13,14^. Longitudinal studies in humans tend to focus on readily accessible organs^15,16^, which may overlook the full spectrum of aging in the body. Moreover, the current predominant focus on the cellular and molecular aspects of aging^17^, while essential, may neglect the intricate interplay between cellular, microanatomical and organ- and organism-level changes throughout the human lifespan. Furthermore, researching aging in humans has the added challenge of distinguishing pathological alterations^18^ from the normative aging process, making it difficult to pinpoint the initiation processes of age-related diseases.

To address these challenges, it is imperative to move beyond a gene, cell, or focus to a holistic view of the aging organism^19^. Deep learning^20^ has greatly advanced our ability to quantify and understand tissue morphological and architectural patterns^21,22^ and detect and classify diseases^23,24^. Here, we employ a novel and systematic approach to probe the interplay between aging and tissue architecture in humans and its connection to pathology, illuminating the complex relationship between tissue-specific aging patterns and their implications across the human body.

### Aging strongly affects healthy tissue architecture in humans

To study the impact of aging on tissue structure in humans, we used large-scale whole-slide histopathological images (WSIs) from 983 individuals in the Genotype-Tissue Expression Project^25^ (GTEx) (**Fig. 1a**). The individuals profiled in this cohort died primarily from accidents, suicide or natural death, with ages ranging from 20-70 years (52.77 mean) (**Fig. S1a**). Tissue was collected under a rapid autopsy protocol from a total of 40 different tissues in 29 organs, making up 25,713 WSIs (**Fig. S1b-e**), which have been reviewed by pathologists and include a description of subclinical levels of pathology (**Fig. S1f-h)**, with demographic, lifestyle and clinical information also available (**Fig. S1i)**.

**Figure 1:**
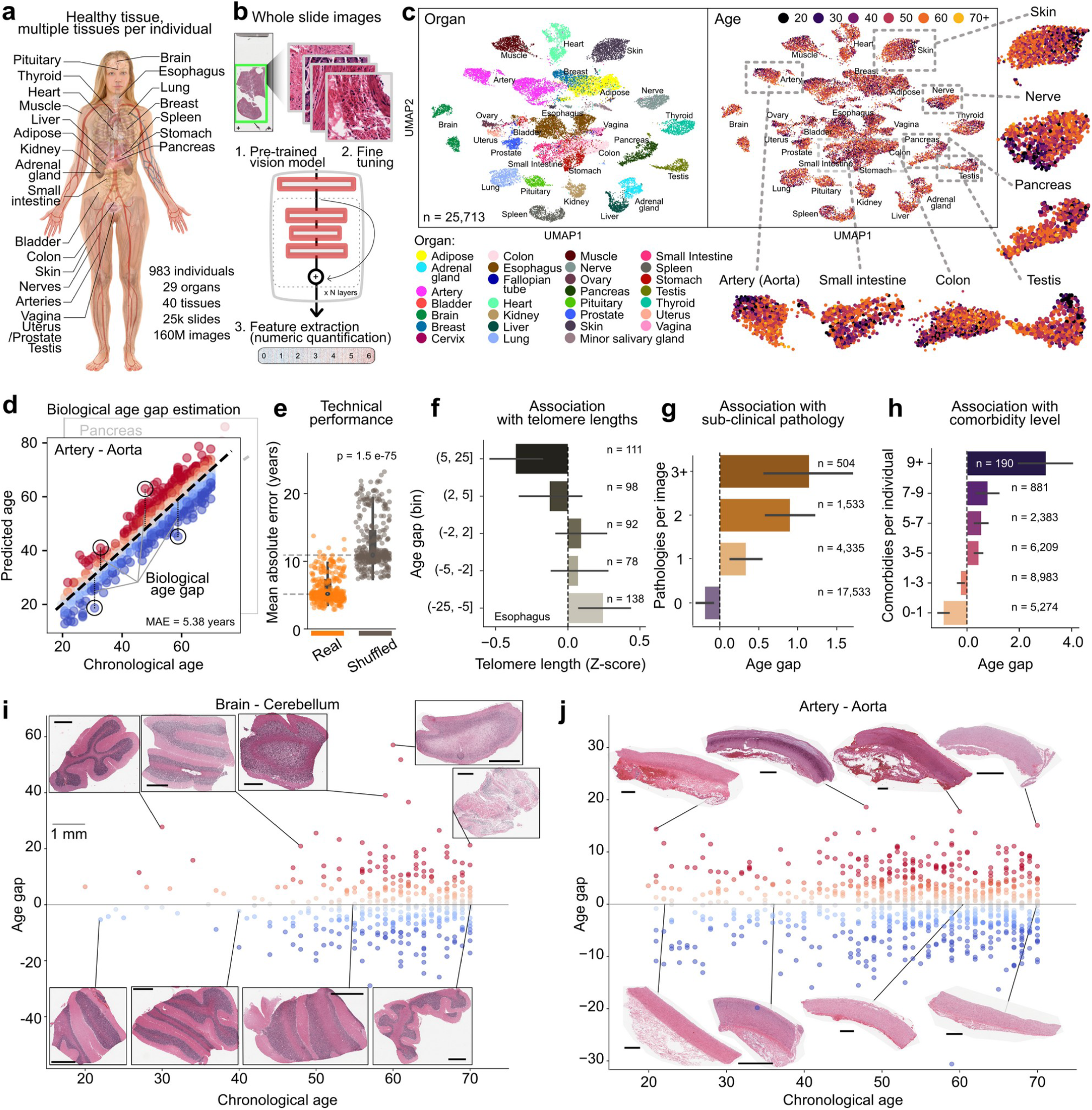
Assessment of biological age and pathological deviations in human tissues via histopathological images. **a)** Overview of the cohort tissue collection approach in the GTEx project, which spans 29 organs and 40 tissues from 983 individuals. **b)** Workflow of image processing from whole-slide histopathological images, illustrating the transformation of raw image data into a structured feature set for quantitative analysis. **c)** UMAP visualization of the processed image data for each WSI, with samples color-coded by organ type (left) and age (right). **d)** Illustration of ‘tissue clocks’ for the aorta to produce estimates of biological age from WSI features. **e)** Assessment of the technical performance of age prediction models using real and shuffled age data. **f)** Relationship between telomere length and the age gap in the esophagus, indicating that greater age gaps are associated with telomere shortening. **g)** Enrichment in the number of subclinical pathologies with increasing age across tissues. **h)** Enrichment in the number of comorbidities of an individual with their predicted age gap. **i-j)** Scatter plot of chronological age and age gap for the cerebellum (i) and aorta (j), illustrating morphological changes associated with chronological and biological age.

Modern AI vision models are powerful extractors of deep features from images that can then be linked to information such as clinical outcomes^21–24^. For our aim, to numerically quantify the morphological features in these WSIs, we first fine-tuned vision models on a set balanced for tissue, sex and age brackets (**Fig. S2a-b**) and then used them to extract a feature vector representing the morphological features of the WSI (**Fig. S2c**). We observed that these features were well reflective of WSI tissue identity even in pretrained models without fine-tuning (**Fig. S2c**), but fine-tuning provided better separation (**Fig. S2d**). More importantly, we found that deep learning morphological features also contained information on individual-level covariates, such as age, even though they were not fine-tuned for that task (**Fig. 1c**, **Fig. S2d-h**). Indeed, we found that age was the strongest factor driving variation across tissues (**Fig. S3**), highlighting the influence of aging on tissue structure throughout the human lifespan.

### Tissue-clocks – estimators of biological age from tissue images predict biological, pathological, and clinical factors associated with age

Building on the observation that age has a marked imprint on tissue structure, we sought to develop ‘tissue-clocks’ tissue image-based predictors of biological age, building on the concept first introduced with DNA methylation^26^. To do that, we trained regression models to predict the chronological age of each individual from the morphological features of the WSI in a cross-validated manner and observed the deviation of the predicted age from the known chronological age: an age gap (Fig. 1d). This gap in DNA methylation data has been associated with several age-related diseases, such as Alzheimer’s^27^, Parkinson’s^28^ and Huntington’s^29^ diseases, amyotrophic lateral sclerosis^30^, and all-cause mortality^31^. We used the model predictions to assess its performance and found an average mean absolute error of 4.88 years across all tissues and a coefficient of determination (R2) of 0.69 (**Fig. S4**, **Table S1**). We also assessed the model robustness by shuffling the ages across individuals and observed no evidence of overfitting; however, 4/40 clocks were underpowered, as fewer than 100 samples were available for these tissues (Fig. 1e, **Fig. S4a-c, Table S1**).

To understand the biological relevance of the tissue-based prediction of age and the value of the inferred age gaps, we leveraged telomere length measurements made from the same tissues and individuals available for 6,197 samples (25.2%)^32^. We observed that as with chronological age, there was an overall decrease in telomere length with biological age (**Fig. S5a-b**). However, this trend was stronger in terms of biological age, as determined through tissue analysis, than in chronological age and was still present in the estimated age gaps (Fig. 1f, **Fig. S5a-e**). This effect was particularly pronounced for tissues such as the esophagus, stomach, kidney, prostate, and pancreas (**Fig. S5c-d**). Furthermore, by leveraging the annotations of subclinical levels of pathology from the same samples, we observed an enrichment of greater age gaps in samples with more pathology (Fig. 1g). This enrichment was concordant across the vast majority of pathologies but, importantly, was greater in tissue-driven estimation of biological age than in chronological age alone (**Fig. S6a-b**), with, for example, both enriched in artery calcification. However, age gaps also recovered enrichment in pathological features not particularly present in chronological age, such as muscle atrophy (**Fig. S6c-d**). Finally, we observed an enrichment in the number of comorbidities an individual has with higher age gaps (Fig. 1h). Confirming the observations that higher age gaps are associated with tissue-specific pathology and comorbidities, we could visually confirm that WSIs with higher age gaps acquired noticeable changes in tissue-specific pathology (Fig. 1i**-j**). For example, independent of chronological age, cerebellum samples with greater age gaps presented discoloration associated with the loss of myelin and ischemic changes (Fig. 1i), whereas in the aorta, samples with higher age gaps presented thickening of the wall and disruption of integrity associated with atherosclerosis and other vascular disorders (Fig. 1j).

### Histological and molecular interpretation of tissue aging

To characterize the histopathological features associated with the aging of tissues more systematically, we leveraged a pretrained self-contrasting multimodal model trained on many pathological images and their natural language descriptions^33^ (Fig. 2a). With this model, we queried the similarity of each image to a set of histological and pathological terms (n = 150), generating a highly interpretable numerical vector to further describe the images. The histological text terms showed high organ and tissue-specific enrichment, with terms such as ‘adipose tissue’ high in breast tissue and visceral and subcutaneous adipose tissue or ‘neurons’ in the brain (Fig. 2b, **Fig. S7a**). We then characterized the association of each term with aging in each organ (**Fig. S7b-c**) and found that terms such as atrophy, microvascular rarefaction and fibrosis increased across multiple organs, whereas the most common reduction in text terms across organs was related to the epithelium and hyperplasia (Fig. 2c**-d**). In addition to these common trends across organs, we also found some more specific trends in certain organs, such as fat infiltration in skeletal muscle (Fig. 2e), which we could visually confirm. Similarly, we also visually confirmed additional associations, such as microvascular rarefaction in the uterus (Fig. 2f) and peripheral nerve atrophy (Fig. 2g). These findings support our deep learning-driven analysis of tissue and enable a histological interpretation of age-associated changes in healthy tissue at a large scale.

**Figure 2:**
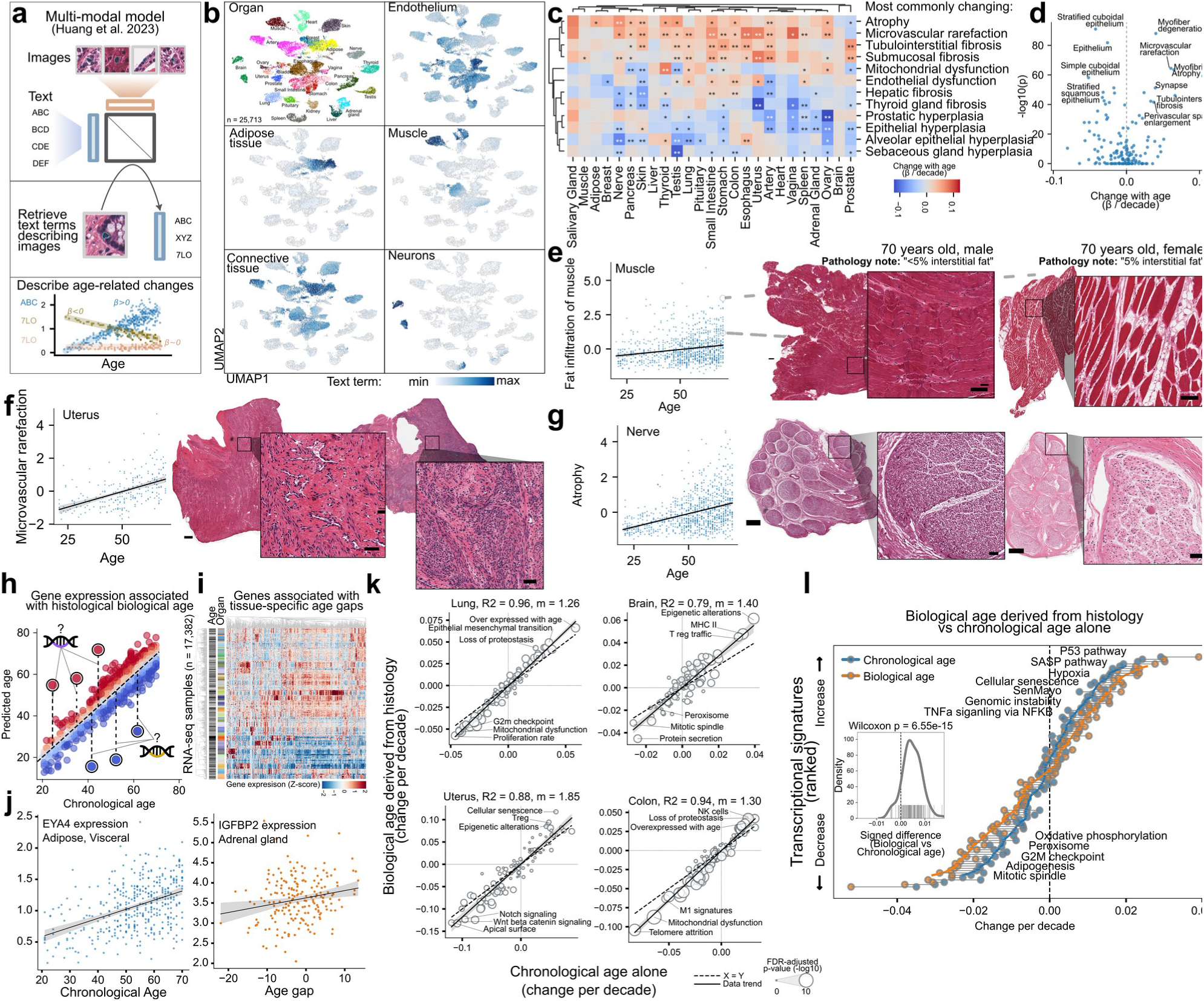
Characterization of age-associated changes in tissue from histological and molecular perspectives. **a)** Schematic representation of the strategy for querying of text-image multimodal pathological models. **b)** Illustration of histological text terms characterizing various organs using the UMAP representation from Fig. 1C. **c)** Heatmap with most commonly changing text terms colored by the magnitude and direction of change for each organ. **d)** Volcano plot on changes in text terms with age across organs. **e-g)** Example of the change in specific text terms with age selected organs (left), with examples on the right (right). Scale bars 500 and 50 microns (inset). **h)** Schematic representation of the approach for the integration of histological and gene expression data to determine gene associations to age gaps. **i)** Comparison across tissues of clock performance (MAE) in biological age prediction, number of genes changing with age, and number of genes associated with histological age gap. **j)** Gene expression for genes associated with age gaps across multiple tissue samples, organized by tissue type and age. **k)** Comparison of changes in transcriptional signatures between chronological age (x-axis) and histologically derived biological age (y-axis). Note the systematic deviation from the x=y dashed line, indicating generally higher changes with biological age. The top and bottom 3 changing terms are identified. **l)** The mean amount of change (x-axis) in each transcriptional signature (y-axis) across organs is illustrated for chronological age (blue) and histologically derived biological age. The distribution of differences between chronological and biological age is displayed as an inset. The sign of the differences were inverted is the mean of the changes was negative.

Building upon these morphological insights, we leveraged the wealth of molecular data from the same samples (**Fig. S9a-b**) to explore the molecular basis of tissue aging.

First, we sought to understand the relationship between estimates of biological age derived from DNA methylation^26,34–38^ and those derived from histological images—tissue clocks (**Fig. S8**). In the GTEx cohort, 987 samples from 8 tissues had available DNA methylation data^39^, which we used to derive a range of biological age estimates via established biological age clocks (**Fig. S8a-b**). We observed that estimates from the best DNA methylation clocks were not as strongly enriched in tissue-specific pathological factors (**Fig. S8c-d**) and that histology- and DNA methylation-derived age gap estimates showed no agreement with each other (0.09 Pearson correlation) (**Fig. S8e-g**).

Second, leveraging over 17,000 matched bulk RNA-seq samples, we investigated how gene expression changes with chronological age and histologically derived biological age. We employed regularized regression models to associate the expression of specific genes with varying degrees of deviation from chronological age in the same samples (Fig. 2h). We observed that many more genes were associated with changes with chronological age than with age gaps (thousands vs. hundreds, **Fig. S9c-e**) but that the degree of association was not necessarily driven by the performance of the tissue clock or sample size (**Fig. S9c**), suggesting that tissue-specific factors may also affect biological age. We found that genes associated with tissue-specific age gaps were overall also tissue-specific (Fig. 2i**, Fig. S9f**) but also included genes not usually expressed in the tissue. For example, we detected the upregulation of EYA transcriptional coactivator and phosphatase 4 (EYA4) in adipose tissue during aging (Fig. 2j), which is usually restricted to the muscle, tongue and brain and was previously associated with hearing loss^40^. Insulin-like growth factor binding protein 2 (IGFBP2), which is usually restricted to the liver, pancreas and stomach and promotes oncogenic processes, was also found to be upregulated in the adrenal gland with increasing age (Fig. 2j). These gene expression profiles offer potential biomarkers for aging and highlight the kind of gene-specific shifts that may contribute to the aging process.

To study how transcriptional pathways and signatures are altered with aging, we leveraged a set of 50 hallmark pathways from MSigDB, as well as 50 aging-related transcriptional signatures, which include processes such as cellular senescence and signatures such as SenMayo, among others. Furthermore, we compared the changes associated with chronological age with those associated with histologically derived biological age to determine the degree to which they provide similar molecular insights into tissue aging. We observed good overall agreement between chronological and histological biological age (mean R^2^ across organs, 0.89), with significantly upregulated pathways most related to apoptosis and inflammation and downregulated pathways most related to metabolism (**Fig. S10b**). However, each organ showed specific changes, likely related to its specific physiological function (Fig. 2k**, Fig. S10c**). For example, epigenetic alterations were among the most upregulated pathways in brain and uterine tissue, whereas in the lung and colon, the pathways associated with the greatest increase in expression were loss of proteostasis and a previously known signature of gene overexpression with age. When closely analyzing these data, we noticed that, despite being similar, estimates of change with biological age consistently presented greater changes than chronological age did, regardless of their up- or downregulation direction (Fig. 2l). Transcriptional signatures such as p53 signaling, cellular senescence and senescence-associated secretory phenotype (SASP), hypoxia, genomic instability, and TNF alpha signaling were upregulated with both chronological and biological age but consistently increased with biological age. Conversely, the mitotic spindle, G2M checkpoint, adipogenesis, peroxisome and oxidative phosphorylation pathways were more downregulated with increasing biological age. On average, the slope between biological and chronological age was 1.26, and the average difference between them for each pathway was significantly different from the expected zero value. Overall, the transcriptional changes associated with histologically derived biological age are similar but recover a greater degree of transcriptional dysregulation than chronological age does, which couldpotentially be used to refine our understanding of the interplay between molecular and histological changes in tissues during aging.

### Synchronicity and deviation in tissue-specific aging clocks

The molecular analysis above suggests that tissue-specific and intrinsic programs condition the process of aging that manifests at the tissue level. To decipher the complex interplay of aging across different tissues, we asked which tissues are most similar in terms of their aging rates throughout the human lifespan. Using a cross-inference approach, where one tissue’s clock was used to predict the age of all others and the process was repeated for every tissue type, we were able to register deviations in predicted age across the population between tissues, which indicated differential aging rates across tissues (Fig. 3a). The resulting matrix of age predictions provided a landscape of relative age acceleration or deceleration across tissues, with some tissues appearing to age faster—predicting others to be years older—while others showed the opposite effect—predicting others to be years younger (**Fig. S11**).

**Figure 3:**
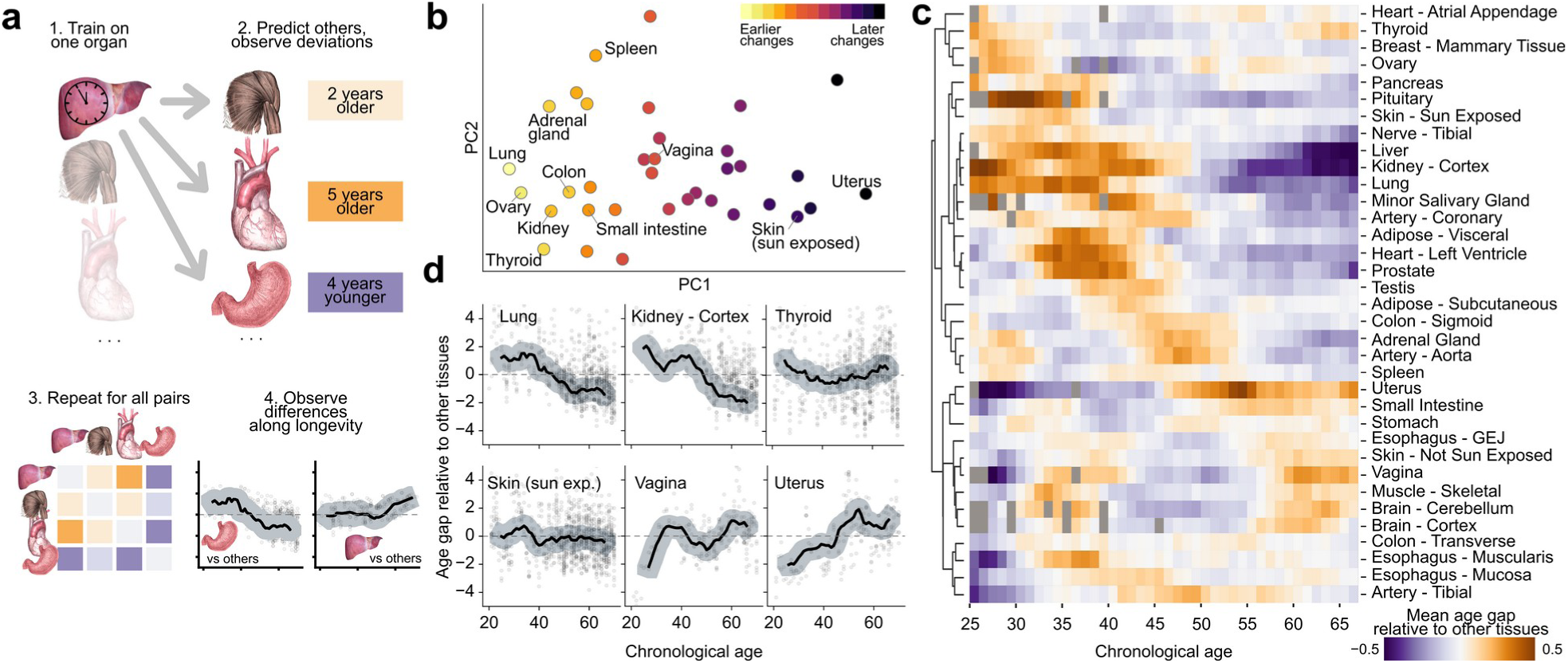
Relative acceleration of tissue-specific aging across the human lifespan. **a)** Schematic representation of the cross-tissue age prediction methodology. **b)** Hierarchy of temporal changes across tissues created through reduced representation of pairwise cross-predictions. **c)** Relative age acceleration of individual tissues with respect to chronological age. The color reflects the degree of relative age acceleration in comparison with other tissues. **d)** Illustration of the rate of age-related changes in selected tissues with different trajectories across the chronological age spectrum.

We then sought to identify a hierarchy of changes across tissues by leveraging dimensionality reduction methods to derive an axis of rates of change between tissues across the human lifespan. Notably, we found that the lung and a group of organs rich in glands, such as the ovary, thyroid, kidney, pancreas and adrenal gland, were among those with overall greater predicted age (Fig. 3b). By stratifying the predicted age gaps and therefore the inferred relative age acceleration rates across the range of human longevity, we noticed a more nuanced view (Fig. 3c), with several organs having non-constant rates of change and peaks of acceleration in specific lifespan regions. In addition to the previously mentioned group of organs with the greatest acceleration of aging in the second to fourth decades of life (20 to 40 years old), we also identified a group of organs (colon, aorta) with a bimodal pattern of age acceleration characterized by higher rates in the third decade of life but also later (45 to 55 years old), as well as a second bimodal group (brain, muscle, vagina, and sun-protected skin) with peaks in the decades of 30 to 40 years and 60 to 70 years of age. The uterus stood out from all other organs, with an inflection point at approximately 45 years of age, and the peak acceleration coincided with the onset of menopause (45 to 55 years of age) (Fig. 3d). Few organs showed a constant rate of aging, which highlights the extremely dynamic nature of the aging process across tissues and suggests that different external factors may affect different tissues at different rates throughout the human lifespan.

### Demographic and clinical factors modulating tissue-specific aging

While the previous analysis highlighted the modulation of the relative speed of aging in different tissues, it does not account for deviations from chronological age that are individual specific. We were interested in quantifying the degree to which aging acceleration in relation to individuals of the same chronological age may be different for different organs within a single person. By leveraging age gaps estimated for various tissues of the same individual (Fig. 4a), we were able to identify several patterns of cross-tissue aging within individuals: consistently below average age gaps across tissues (‘resilient agers’); consistently average age gaps (‘average agers’); single- and multiple-tissue agers – one or a few tissues with greater age than others; and a group of individuals whose most tissues were consistently predicted with greater age gaps (‘systemic agers’). To visualize the pattern of aging across many individuals, we focused on a set of 303 individuals with greater than average age gaps in at least one tissue and identified various groups of individuals on the basis of their internal organ synchronicity in aging rates (Fig. 4b). We observed that most individuals with at least one high age gap tended to have several organs with large age gaps (Fig. 4b, pan-tissue cluster). However, we detected several individuals with one particularly prominently aged organ, such as the heart, lung or stomach (Fig. 4c).

**Figure 4:**
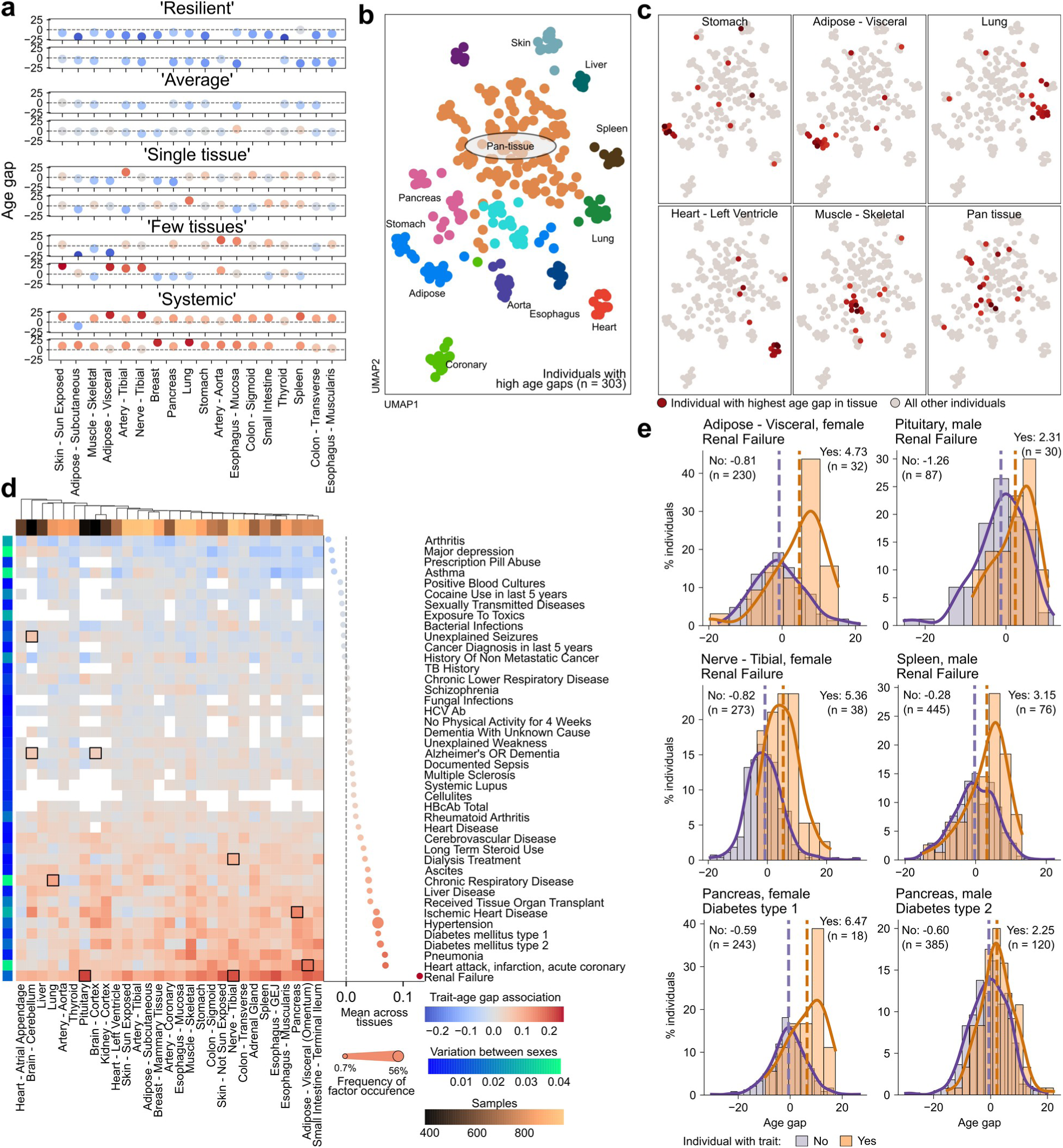
Diversity of aging within and between individuals. **a)** Inferred age gaps across tissues in example individuals depending on the pattern of aging across tissues. **B-c**) UMAP plot demonstrating the variation in age gaps across tissues for each individual. Individuals are colored according to the pattern of inter-tissue variation during aging (b) or according to the tissue with the greatest age gap. **d**) Association of various demographic, clinical, and lifestyle factors with tissue-specific age differences. The color denotes the log hazard ratio. Boxes highlight known associations or are highlighted in panel e) or the text. **e**) Histograms comparing inferred tissue-specific age gaps depending on clinical factors. Individuals were split by whether they were annotated with a factor, and the mean age gap and size of each group are shown.

Interestingly, none of these organs were part of the organs that were mostly aged ‘late’ (Fig. 3c), which may suggest that early events causing accelerated aging may have a cumulative effect throughout life. To identify factors potentially underlying a specific deviation in the aging of one organ in comparison with others among the 983 individuals, we devised a strategy to associate age gaps for each tissue with demographic, clinical, and lifestyle information. These included 184 variables, from which we identified 80 that were recurrent enough to assess in light of the tissue-specific age gaps (**Fig. S1i**). We discovered that, by and large, most factors tended to affect several or most tissues (Fig. 4d**, Fig. S12a-b**) but also revealed several associations that were tissue- and sex-specific. One of the most prominent observations was the association of renal failure with accelerated aging across several tissues (Fig. 4d). Interestingly, the strongest association was not found in the kidney but was detected in tissues such as adipose tissue and the pituitary, spleen and tibial nerves (Fig. 4e). In females, the association between tibial nerve aging and kidney failure could be related to the fact that more than 90% of chronic kidney disease patients who undergo dialysis suffer from peripheral neuropathy^41^. With respect to variations across tissues, we confirmed several known associations, such as chronic respiratory disease affecting most prominently deviations of lung tissue biological age from chronological age^42^ (Fig. 4d), type II diabetes associated with pancreas aging^43^ (Fig. 4d**-e**), heart disease associated with adipose tissue aging^44^ (Fig. 4d), and the effects of steroid use on the spleen^45^ (Fig. 4d). However, we also found other less well-known associations that have not yet been linked to an explicit morphological manifestation of the aging process, such as those associated with higher rates of aging in the cerebellum with unexplained seizures^46^ (Fig. 4d) or accelerated prostate aging and hypertension in men (**Fig. S12b**), even though there are reports that link hypertension with prostate hyperplasia^47,48^.

### Blood-based prediction of tissue-specific age gaps

Given that gene expression could be associated with tissue-specific biological age derived from images (Fig. 2) and that aging is reflected in a systemic manner in many individuals (Fig. 5), we aimed to develop a predictor of tissue-specific biological age (driven by morphological features in tissue) from gene expression samples of blood in the GTEx cohort (Fig. 5a). By linking blood-based gene expression profiles with the histology-derived tissue-specific age gaps of the same individuals, we were able to derive blood-based predictors of tissue age gaps across tissues (Fig. 5b). Among all the predictors, we observed good performance in the prediction of tissue-specific age gaps in the GI tract and the spleen (Fig. 5b). However, the best performance was observed for a predictor of the mean number of tissue gaps in each individual (Fig. 5B, ‘systemic’), which may suggest that blood gene expression has the greatest deviation from random in systemic tissue aging signals, reflecting an integration of the aging process of several tissues and organs. Finally, we observed that the predicted age gaps from blood were negatively associated with the length of telomeres in tissue and highly enriched at the level of pathologies across all tissues of an individual (Fig. 5c**, Fig. S13**). We examined the coefficients of the genes underlying the predictive process and found that they were stable and well distributed indicating robustness (**Fig. S14a**). Furthermore, we took further advantage of the interpretability of linear models found that overall, each tissue-specific predictor had a specific set of unique genes and that genes positively associated with age gaps in one tissue and negatively associated with another (**Fig. S14b-c**). Interestingly, we found several genes not known to be expressed in blood to be among the top predictors of tissue-specific age gaps (**Fig. S14c**). For example, junction plakoglobin (JUP) is expressed in glandular and epithelial cells and has the strongest association with the small intestine, skin, and adipose tissue, as well as with systemic predictors. Similarly, the expression of keratin 5 in the blood, which is traditionally a marker of basal epithelial cells, had the strongest association with all genes predicting pancreas age gaps.

**Figure 5:**
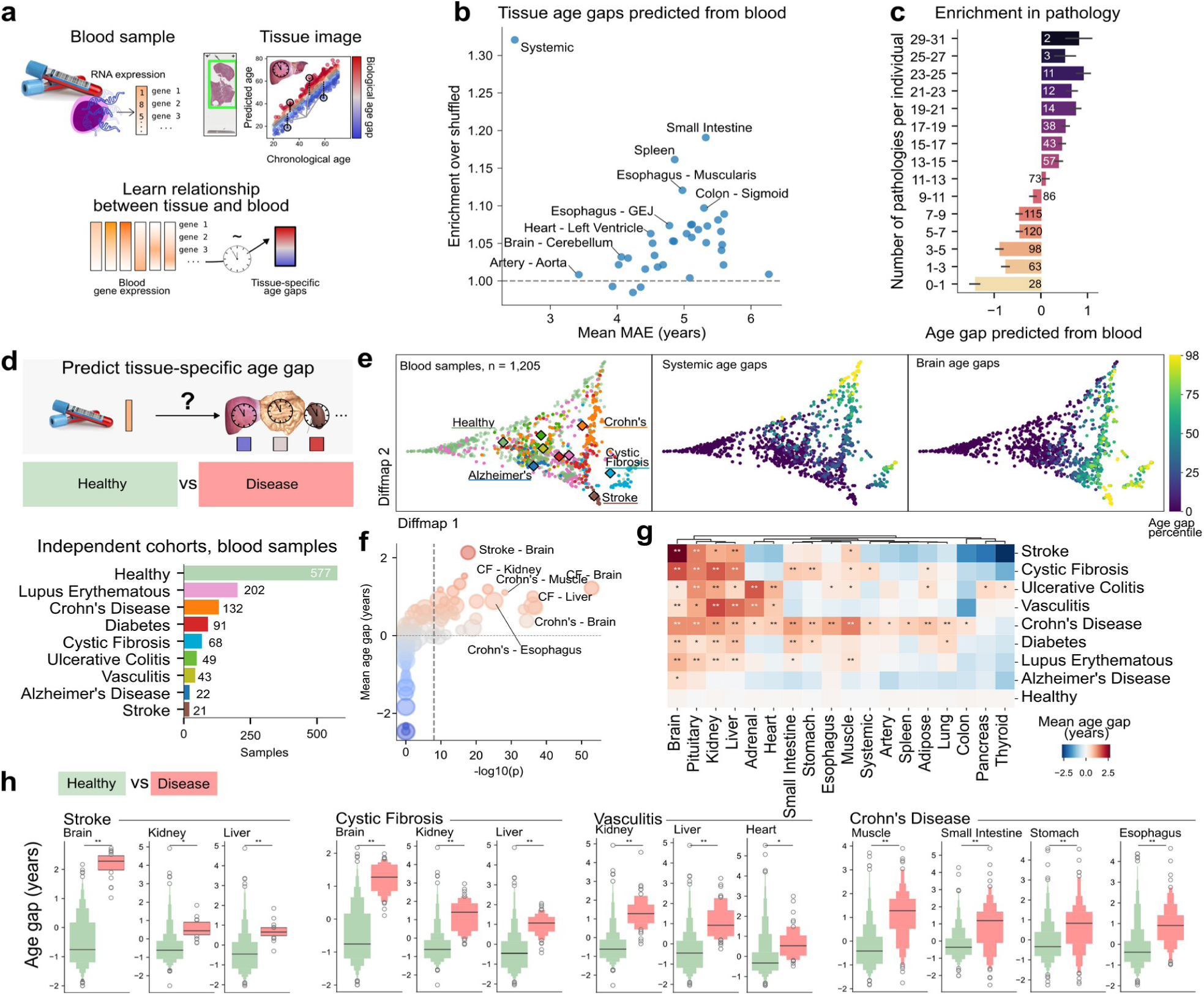
Prediction and validation of tissue age gaps from blood gene expression. **a**) Overview of the strategy combining image-based prediction of biological age gaps from WSIs with blood gene expression to predict tissue-specific age gaps from blood. **b**) Scatter plot showing the enrichment of performance in blood-based prediction of tissue age gaps for various tissues and for the mean of age gaps across tissues for each individual. **c)** Enrichment in the annotated tissue pathologies in age gaps derived from blood prediction of an individual’s systemic tissue age gap. **d)** Overview of the composition of the independent cohorts used to validate the blood-based tissue-specific clocks. **e)** Diffusion map representation of the age gaps estimated from blood for the 1205 samples revealing a landscape of disease-. **f)** Volcano plot of the association of age gaps with pairs of diseases and organs. **g)** Heatmap of mean age gaps for each disease in each organ. Significant associations are highlighted with asterisks. **h)** Examples of significant associations from f) and g) found between disease and healthy groups. * p < 1e-3; ** p < 1e-8, T-test, BH-FDR adj.

To validate the blood-based tissue-specific predictors and assess whether they hold insights into affected organs during disease, we leveraged the large-scale repository of harmonized gene expression and sample metadata ARCHS4^49^ to find blood samples of healthy individuals (n = 577) or with one of seven chronic diseases (n = 628): systemic lupus erythematous, Crohn’s disease, diabetes, cystic fibrosis, ulcerative colitis, vasculitis, Alzheimer’s disease (Fig. 5d). As a control, we also included stroke patients representing an acute medical event affecting specifically the brain. We then predicted tissue-specific age gaps based on the coefficients derived from the GTEx cohort, and observed that while there was no relationship with sample size, they were distinct for each disease group (**Fig. S15a-c**). Joint analysis of age gap predictions across all organs revealed that samples were distributed along two axes, with disease entities forming distinct zones ( Fig. 5e and **Fig. S15d-e**). One axis showed a transition from healthy samples with near-zero age gaps to samples from various diseases with higher gaps, whereas the second axis reflected increasing age gaps in multiple organs, particularly in the heart, adipose tissue, and gastrointestinal tract. Samples from stroke and cystic fibrosis patients had especially high age gaps in the brain and kidney, respectively (Fig. 5e). Statistical testing confirmed significant associations between diseases and elevated tissue-specific age gaps (Fig. 5f**-g** **and Fig. S15f**). Notably, stroke patients showed the most pronounced age gap deviation specifically in the brain (Fig. 5f**-h**), in alignment with the affected tissue in this acute event. Moreover all seven chronic diseases were significantly associated with tissue-specific accelerated aging in at least one organ (Fig. 5g). For example, Crohn’s disease was associated with increased predicted age gaps in the gastrointestinal tract (esophagus, stomach, small intestine, colon), whereas vasculitis exhibited higher age gaps in the kidney, liver, and heart (Fig. 5h) which are highly vascularized organs and primary targets of inflammatory processes in this disease. In some diseases, fewer and milder but specific associations were detected; for example, the brain was the only organ with significantly higher age gaps in Alzheimer’s disease (Fig. 5f**-g**). Interestingly, we also found associations of age gaps with organs which are not the primary site of disease action. These associations could be revealing of how treatment of a primary condition may be affecting other organs^50^ as detected by significantly higher age gaps for stroke patients in kidney and liver, or of secondary effects of disease such as changes in the brain due to chronic hypoxia in cystic fibrosis^51^, for which we also detected significantly higher age gaps in comparison to healthy controls (Fig. 5h). Finally, to assess the potential clinical utility of blood-based tissue-specific age gaps in detecting disease, we evaluated thresholded age gaps in their performance (**Fig. S15g**). The models demonstrated moderate to strong classification performance as assessed by the area under the receiver operating characteristic curve, while positive predictive values above 0.3 in several cases suggest potential applicability for population-level screening.

Overall, this approach demonstrates the feasibility of inferring tissue-specific biological age and disease-related aging signatures from minimally invasive blood samples, underscoring its potential utility in early disease detection and monitoring aging in clinical settings.

## Discussion

This study presents a comprehensive analysis of human aging through tissue histology, highlighting the complexities of tissue-specific age variation across tissues and within individuals. The GTEx dataset provided an unprecedented view of histopathological hallmarks of aging in both healthy and subclinically affected tissues, enabling correlations with molecular, demographic, and clinical factors.

The morphological assessment of tissue underscored the profound impact of aging on tissue architecture. Despite the commonality of aging as a universal process, we revealed a multitude of factors modulating it and a remarkable variation in the rates of aging across tissues during the human lifespan. The development of ‘tissue-clocks’ stands, therefore, as a pivotal advance, following the paradigm first established with DNA methylation studies^26,34–38^ but bringing a highly tissue- and organ-specific context that is deeply connected to physiological fitness.

Leveraging the largest dataset of paired histological images, RNA-seq and DNA methylation data from non-cancerous tissue, and multimodal deep learning models, we were able to systematically characterize the histological features of tissue aging and identify loss of epithelial proliferation and increase in tissue atrophy, microvascular rarefaction and fibrosis as some of the most common aging features across organs. The observation that changes in pathways and transcriptional signatures were more strongly recovered with histologically derived biological age than chronological age suggests that histological analysis captures a more pronounced molecular aging signal, potentially reflecting the cumulative burden of microanatomical alterations to tissue that are not fully accounted for by intrinsic cellular changes alone. We observed agreement between the age estimates derived from DNA methylation and histology only in biological age, although both methods resulted in low errors and association of age gaps with pathological features. Moreover, it is noteworthy that histological measures of biological age and its gap with chronological age align with telomere attrition rates, particularly in tissues of the gastrointestinal tract, lung, pancreas, and kidney. While agreement between epigenetic aging clocks can be fairly low^52^, the lack of agreement we observed between tissue clocks and DNA methylation estimates could be because DNA methylation clocks leverage the accumulation of stochastic changes in DNA^53^, whereas tissue morphology is conditioned by physiological and external factors that diverge from those driving stochastic DNA changes. Giving credence to that, the analysis of gene expression suggested that some molecular pathways are driven more by changes associated with chronological age and that histological age gaps can be associated with pathways that exacerbate or dampen these effects in a tissue-specific manner. This finding also suggests that aging is not merely a chronological countdown but a complex process with malleable pathways and is permissive of intervention.

With the increasing efforts of large-scale histopathological^54,55^ repositories providing an avenue for the large-scale validation of tissue-clocks, it is conceivable that they may be applicable in tissues with routine sampling, such as during colonoscopy. However, physiologically relevant, tissue image-based clocks have the disadvantage of requiring the invasive procurement of tissue. In the search for accurate biomarkers of aging, noninvasive or minimally invasive methods such as thermal facial imaging^56^ or plasma proteomics are desirable. The latter have recently shown the ability to make organ-specific inferences by making assumptions on the origin of circulating proteins in plasma^57–59^. In our work, we utilized the observed changes in tissue morphology to train a blood-based predictor of tissue-specific aging rates, which was enriched in tissue-specific pathologies and which we were able to validate in independent cohorts of patients with eight different diseases. This analysis, uncovered specific signal for affected organs in both acute events (stroke) but even in remarkably heterogeneous chronic diseases such as significantly higher age gaps for brain in Alzheimer’s disease patients. To assess the clinical utility of blood-based predictors, we evaluated their classification performance, achieving moderate to strong predictive power with positive predictive values (PPV) exceeding 0.3 in some cases—a threshold relevant for population-level screening. However, the robustness of these models under data distribution shifts remains to be further assessed. Interestingly, significant associations of tissue-specific age gaps in organs beyond primary disease sites, such as the pervasive accelerated aging of kidney and liver across multiple chronic diseases in comparison with healthy individuals. The ability to detect both organ-specific and systemic aging patterns from a minimally invasive blood test underscores the potential of this approach not only as a tool for early disease detection in a future prognostic marker, but also for disease and treatment monitoring.

However, in reflecting on the potential of our research, we acknowledge several crucial limitations that accompany our findings: 1) A notable aspect is the sex imbalance within the GTEx cohort (1:2 female:male ratio). This disproportion is reflected in the performance of aging clocks, with 3/4 (from a total of 40) underperforming clocks being from female reproductive system tissues; 2) The use of postmortem samples poses a challenge owing to the variable onset of tissue autolysis between death and the start of autopsy. Due to the excellent annotation of this factor, we have adjusted for this to reduce its potential impact, but we cannot exclude that that has reduced the power for discovery in some tissues; 3) Due to the use of cross-sectional data collected post-mortem, our analysis is not well powered to address questions with causal relationships, but currently recovers existing correlations. In future endeavors, combining genetic data with approaches such as genome-wide association studies (GWAS) and Mendelian randomization, similar to studies conducted in larger cohorts without tissue-level data^60^, may provide avenues to infer causal relationships.

Despite these limitations, our study stands as a testament to the potential of tissue analysis in aging research and supports the view of aging as a dynamic interplay of molecular, cellular, and histological changes that are intricately influenced by an individual’s life trajectory^3,19^. Furthermore, by connecting quantitative tissue-derived aging phenotypes to molecular data from blood, we provide a window to fingerprint the physiological state of multiple organs from a clinically amenable blood sample in health and in age-associated diseases.

## Methods

### Data reporting

No statistical methods were used to predetermine sample size due to the predetermined sample availability of the GTEx Project^25^. 3 samples were excluded from the study as they were annotated with the wrong tissue label.

### Data acquisition

Whole-slide images were downloaded from the GTEx public portal^25^. All the slides were acquired at 20X magnification (∼0.5 microns per pixel) in the same scanner type (Aperio svs format). Demographic data (including the exact chronological age of the individuals), lifestyle information, clinical information for the participants, and gene expression data were downloaded from dbGAP since they are under protected access.

### Whole-slide image processing

Whole-slide images were segmented into a foreground tissue area and a background area via computer vision algorithms as previously described^61^. We adapted the code from this work, repurposing it into a general WSI package that is independent of the model implemented and that we now open source (see **Code availability**). Briefly, a WSI thumbnail’s signal was converted to HED color space, Otsu thresholded, dilated, and small objects (<500 pixels) were removed. To produce a tissue mask, the holes were filled, and a separate mask for the holes (>0.5 and <50 pixels) was used. Contours were stored in h5 format, and slides were tiled in the tissue area with square patches of 224 pixels.

### Vision model fine-tuning

To capture histological features better, we fine-tuned a variety of vision model architectures in a tissue classification task. Importantly, we created a balanced dataset comprising equal numbers of slides of different tissues, age brackets and sexes. We built upon well-established and pretrained models (AlexNet, VGG16, ResNet50, ResNet152, ResNet50, ConvNext tiny and ConvNext base), with a classifier head corresponding to the tissue classes. To train the new classifier head layer, we first froze all the layers of the pretrained model and trained one epoch, followed by up to 100 epochs of fine-tuning the whole model via the AdamW optimizer with cosine annealing. We started with an empirically discovered learning rate by measuring the loss at incremental steps each step and choosing 1/10 of the minimum (*learner.lr_find*). Model fine-tuning was performed with PyTorch and Fast.ai either on TPUs through Google Collab or on a workstation with two NVIDIA RTX A6000 GPUs.

### Feature extraction and unsupervised analysis

Inference was run on an HPC cluster using CPUs for all tiles in all slides at the 3 levels of magnification (∼480 million tiles). We worked with three different tile widths (224, 448, and 896 pixels) but centered at the same location, such that the tile centroids across different widths are matched and represent different, progressively wider views of the same tissue location. Features were extracted with a variety of models (see section above), and for downstream analysis, feature values were aggregated across tiles by means and concatenated across different tile widths to generate a single feature vector for each WSI. We used AnnData and Scanpy^62^ to perform dimensionality reduction with principal component analysis (PCA) to compute neighbors and compute UMAP for visualization, each step with default parameters (PCA: 50 PCs, neighbor graph: 15 neighbors on the basis of PCA representation, UMAP: 0.5 min_dist, 1.0 spread).

### Analysis of total variance explained

To provide a global measure of the importance of various factors in explaining the observed range of morphological heterogeneity across tissue samples, we employed linear models that use those factors to predict the variability of samples in the principal components. We collected 86 variables related to demography, lifestyle, serology, morbidity, and circumstances of death in each patient. Regression was based on an ordinary least squares regression model (statsmodels^63^) and was fit for each tissue and principal component at a time. The full model was fit first, and models missing each of the 86 variables (leave one out) were fit subsequently and compared to the full model on the basis of the adjusted coefficient of determination (R^2^). These values were weighted by the variance ratio of each principal component in the data and summed across components to generate a value of total variance explained for the factor.

### Tissue-clocks and estimation of age gaps

We employed linear models implemented in scikit-learn^64^, Linear Regression, Ridge, RidgeCV, RandomForestRegressor using GroupKFold (k = 5) cross validation, where the group was an individual, to ensure that the training and validation splits did not contain the same individual. Models were trained for each tissue separately using the features from the vision models as inputs, age as the target, and sex, cohort type and minutes of ischemic time as covariates. To assess potential overfitting, we also fit the models via shuffled age labels. We computed the coefficient of determination (R^2^) and mean absolute error as metrics. Adjustment for regression to the mean was performed by regressing out the coefficient of age from the observed residuals as previously described^16^. All the results reported downstream were obtained for tissue clocks trained with a Ridge model, as it exhibited the smallest mean absolute error, and was fast.

The ‘Bladder’, ‘Cervix - Ectocervix’, ‘Cervix - Endocervix’, ‘Fallopian Tube’, and ‘Kidney - Medulla’ tissues all had fewer than 100 samples available and overall had poor performance (e.g., mean absolute error >9 years) and were excluded from further analysis. We also noticed that 3 samples had extreme age gap values (’GTEX-1GMR2-0426’, ‘GTEX-1S82U-0426’, ‘GTEX-11ZTS-0426’), which, upon inspection were annotated with the wrong tissue type.

### Telomere lengths and annotated levels of tissue pathology

We leveraged the telomere quantity index (TQI) values measured for tissue blocks of the same tissues and individuals in the GTEx cohort^32^. We Z scored these values per tissue to account for intertissue variability in the mean values. Only tissues with at least 100 paired whole-slide images and telomere length samples were used for analysis.

We also leveraged the pathological notes available in the GTEx cohort, in particular the discretized set of 57 text categories describing the whole-slide images, which comprise 11,016 instances of a term annotating an image. For the purpose of associating particular pathological annotations with histological age gaps, we excluded ‘clean_specimens’, ‘no_abnormalities’ and ‘tma’. For pathologies common across more than one tissue, we also derived an overall ‘body pathology burden’, which is a composite measure of the number of pathologies annotated in all tissues of an individual.

### Text term characterization of age gaps

We used the PLIP model^33^ to embed 512 pixel-wide (at 20X or ∼0.5 microns per pixel) tiles from every slide in the GTEx project and used them to query the similarity to a set of histological and pathological terms. The terms were selected to represent equal parts of the histological components and features of the various tissues under study (e.g., epithelium, muscle, neurons, adipose tissue, and myofiber degeneration). The values were aggregated by means per slide, and associations with histological age gaps were derived via regularized linear regression (Ridge).

### DNA methylation clocks

We used the PyAging package^65^ to predict the biological age of the DNA methylation samples matched to the histological images of the tissue. DNA methylation data^39^ were measured via the Infinium HumanMethylationEPIC bead chip, which measures approximately 850,000 CpG sites. We then derived age gaps by comparing them to the real chronological age of the samples and compared both the predicted values and the DNA methylation-derived age gap values to the histological predictions via Pearson’s r.

### Gene expression analysis

We employed bulk gene expression data from the GTEx project that are matched to the same tissues as the histopathology images. We converted gene-level counts to log counts per million (log cpm), and any technical replicates were aggregated by means. To explain both longevity and tissue-specific rates of age acceleration (age gaps) with gene expression of the corresponding tissues, we employed ridge regression using, as previously described, sex, cohort, and minutes of ischemic time as covariates. Genes with an absolute coefficient value above 0.005 (>5% change/decade) were selected. Finally, we used Enrichr^66^ through the gseapy package^67^ for gene set enrichment with the MSigDB database separately for the up- and downregulated gene sets.

Similarly, we also used the 50 MSigDb pathways together with a set of 50 age-related signatures previously compiled (available at https://github.com/WJPina/HUSI/) and quantified them in the transcriptomes via the Scanpy function sc.tl.score_genes. This allowed us to obtain a reduced set of broadly and specifically relevant transcriptional features for each sample. We then calculated their association with chronological age or histologically derived biological age per organ and compared the coefficients between the two per-organ as well as the mean across organs and assessed whether their signed deviation differed via a Wilcoxon test.

### Cross prediction of biological age across tissues

The prediction of biological age across tissues (cross prediction) was performed as described above, using sex, cohort type and minutes of ischemic time as covariates and a Ridge model. To visualize the overall trend in the relative differences between models, we performed principal component analysis on the residuals and aggregated values via an exponentially weighted window of 10 years across the range of chronological age per tissue and aggregated the values via a rolling-centered 10-year window. The clustered heatmap visualizing the relative differences (Fig. 3c) was calculated with cosine distance and average linkage.

### Association of tissue-specific age gaps with individual factors

To visualize individuals in groups with differential patterns of biological aging across the various tissues profiled per individual (Fig. 4b), we selected the tissues with abnormally large age gaps (>=3 standard deviations), set the remaining tissues to zero, and performed principal component analysis (PCA), computed neighbors, and a UMAP as described previously.

To statistically associate individual age gaps with individual-specific factors, we fitted regularized linear models (Ridge) per tissue and sex with factors explaining tissue-specific age gaps. To avoid collinearity, we excluded the following variables: “Abnormal Wbc”, “Drugs For Non Medical Use In 5y”, “HBcAb IgM”, “HIV 1 NAT”, “HIV I II Ab”, “HIV I II Plus O Antibody”, “Nephritis, Nephrotic Syndrome and/or Nephrosis”, “Night Sweats”, “Open Wounds”, “Received Human Growth Hormone”, and “Tattoos Done In 12 m”. Furthermore, we retained only factors with more than 3 individuals per tissue and sex.

### Prediction of age gaps from blood gene expression

The predictions of biological age for multiple tissues we developed are rich, tissue-specific phenotypes relevant to the physiological state of the donor during age and disease. We reasoned that those same age-associated phenotypes could be predicted from blood profiles in order to detect disease states specific to certain organs. To enable that vision, we employed bulk gene expression data from the GTEx project that are matched to the same tissues as the histopathology images. Blood gene expression profiles were log transformed and converted to counts per million (log cpm). We filtered lowly expressed genes by removing genes with a mean log(cpm) <1 (11,859 genes left) and aggregated the technical replicates by means. Since we observed an effect of the aging process on gene expression in general, we opted by first regressing out the effect of age from the blood gene expression profiles via regression. Then, we fit regularized regression models to predict tissue-specific age gaps from blood gene expression via Ridge models with two cross validation loops (RidgeCV): one external Kfold (k = 5) that separates different groups of individuals and where standard scaling is applied and one internal Kfold (k = 5) for alpha hyperparameter optimization (chosen from 10^-1^ to 10^7^ in 20 uniform steps). This process was performed independently for each tissue, and for the mean age gap across the tissues of an individual (systemic clock).

For validation, we obtained a comprehensive set of bulk gene expression profiles from samples of peripheral blood mononuclear cells, leveraging the ARCHS4^49^ database, version 2.2. Gene expression counts were log transformed, normalized in relation to the total, standardized and scaled for each gene, and the coefficients derived for each predictor in the GTEx cohort were applied in a linear model to make age gap predictions for the bulk RNA-seq samples. The primary metric selected to evaluate performance between each disease and healthy samples was a two-tailed t-test with Benjamini-Hochberg false discovery rate correction. This was employed due to the fact that our regression models output continuous values (age gap) which are Gaussian and centered at zero – as those were learned from the residuals of our tissue-clocks models. We also evaluated the utility of the blood-based predictors in a binary setting by thresholding the age gaps at 1, and calculating the area under the receiver operator curve (ROC AUC) and the positive predictive value/precision at separating healthy controls and disease.

## Data availability

GTEx whole slide images, RNA expression and DNA methylation data are available from its portal (https://gtexportal.org). The phenotypic data for the individuals are available from the dbGaP platform under accession phs000424.v9.p2. We make available the balanced datasets used to fine tune vision models at the following publicly available repository: https://doi.org/10.5281/zenodo.13330659. Data from the validation cohorts are available from GEO with accessions GSE66573, GSE72420, GSE72509, GSE84891, GSE89403, GSE102114, GSE109313, GSE110685, GSE111459, GSE112087, GSE113386, GSE120691, GSE123658, GSE123786/7, GSE124548, GSE125512, GSE129752, GSE131475, GSE134080, GSE136053, GSE136371, GSE143507, GSE147931/33, GSE151282, GSE159121, GSE160268, GSE161196, GSE161731, GSE162828, GSE169687, GSE171770, GSE174056, GSE184050, GSE184876/7, GSE185263, GSE186593, GSE187429, GSE199819, GSE206250, GSE206648, GSE208405, GSE212610, GSE222408.

## Code availability

The source code is provided in the Supplementary Information files for reviewers and will be publicly available at the GitHub repository https://github.com/RendeiroLab/agingpath. The package for the analysis of WSIs is available open source at https://github.com/RendeiroLab/wsi, which is based on the “CLAM” repository from the Mahmood lab^61^.

## Acknowledgments

This research was funded by Angelini Ventures S.p.A. Rome, Italy. The Genotype-Tissue Expression (GTEx) Project was supported by the Common Fund of the Office of the Director of the National Institutes of Health, and by NCI, NHGRI, NHLBI, NIDA, NIMH, and NINDS. The datasets used for the analyses described in this manuscript were obtained from dbGaP at http://www.ncbi.nlm.nih.gov/gap through dbGaP accession number phs000424.v9.p2.c1. This work was supported with TPUs from Google’s TPU Research Cloud (TRC). We thank Mikael Häggström for creating the original image used in Fig. 1a and providing it in the public domain (CC0 license). We thank Laura de Rooij and Giulio Superti-Furga for helpful comments on the manuscript text.

## Conflicts of interest

E. A., I. B., Y.Z, and A.F.R. declare competing financial interests in the form of a pending patent application on work developed in this manuscript.

## Notes

### Summary of Updates

Addition of independent validation cohorts in Figure 5.

