## Supplementary Figures for "Histological aging signatures enable tissue-specific disease prediction from blood"

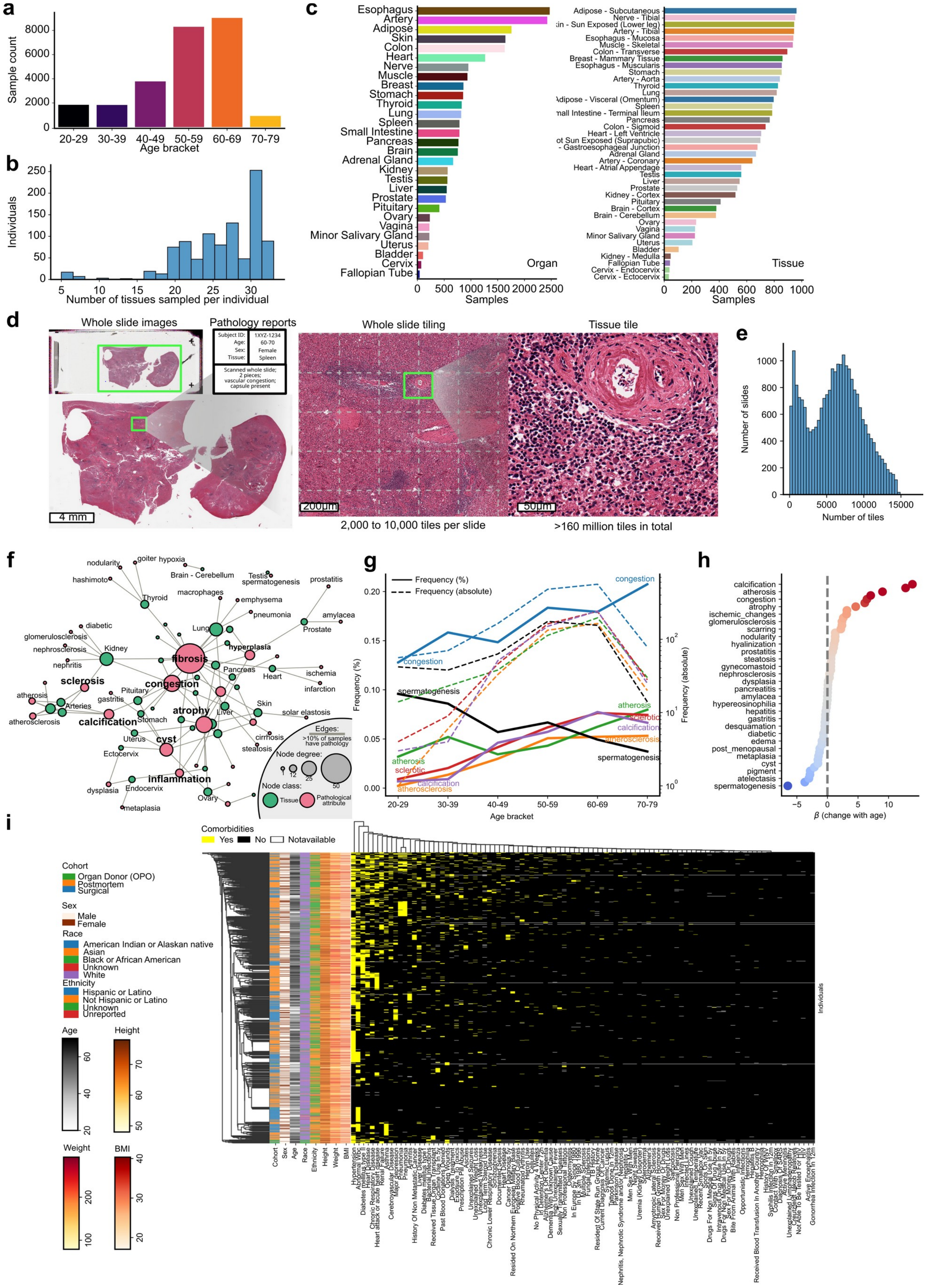

**Fig. S1**: Histopathological, demographic, and clinical data in the GTEx project. **a**) Distribution of sample along age brackets. **b)** Distribution of the number of tissues sampled per individual. **c)** Number of samples per organ (left) or tissue (right). **d)** Example histopathological image highlighting its multiple scales (whole tissue, microanatomical and single-cell). **e)** Distribution of image tiles per WSI in the dataset. **f)** Network illustrating co-occurrence of pathological annotations in tissues. Green nodes are tissues and pink pathological categories with edges connecting them if at least 10% of samples of one tissue are annotated with that pathology. **g)** Relative and absolute frequency of pathologies per age bracket. **h)** Inferred rate of change in pathological term incidence dependent on age. **i)** Heatmap illustrating the frequency and co-occurrence of demographic, clinical, and behavioral factors on the cohort.

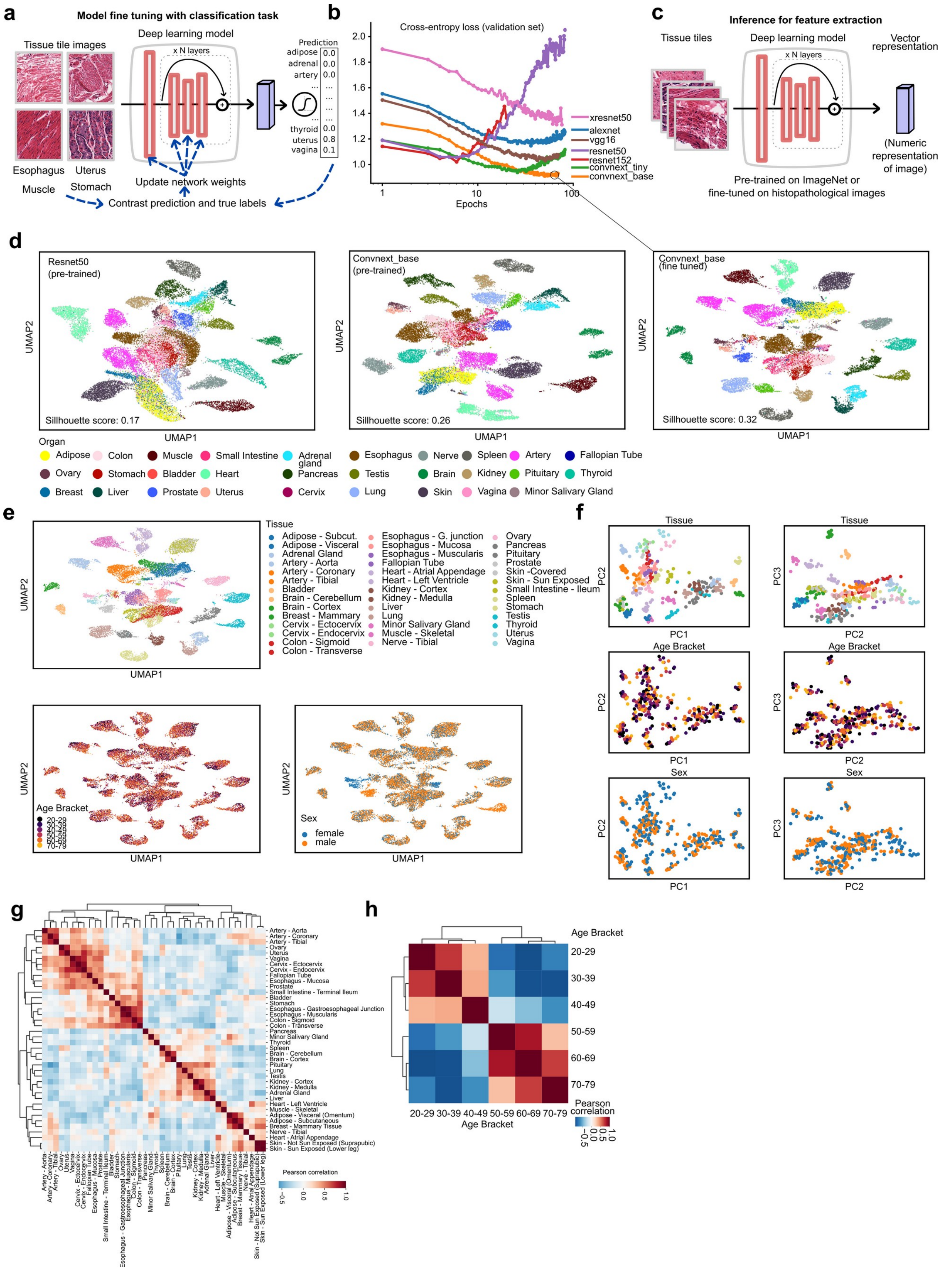


**Fig. S2**: Vision models for the analysis of histopathological images across tissues. **a**) Strategy for the supervised fine-tuning of vision models by classifying tissue types. Datasets were balanced per tissue, age bracket and sex. **b)** Performance of various model architectures during fine-tuning. **c)** Strategy to use vision models for feature extraction. **d)** UMAP plot of all WSIs comparing 3 models: a pre-trained Resnet50 and Convnext base, and a fine-tuned Convnext base. WSIs were colored by organ. **e)** UMAP for a fine-tuned Convnext base model as in d) but with WSIs colored by tissue, age bracket and sex. **f)** PCA plots for the fine-tuned Convnext base model with WSI features aggregated by tissue, age and sex. **g-h)** Pairwise correlation plots for the WSI features aggregated by tissue (g), or age bracket (h).

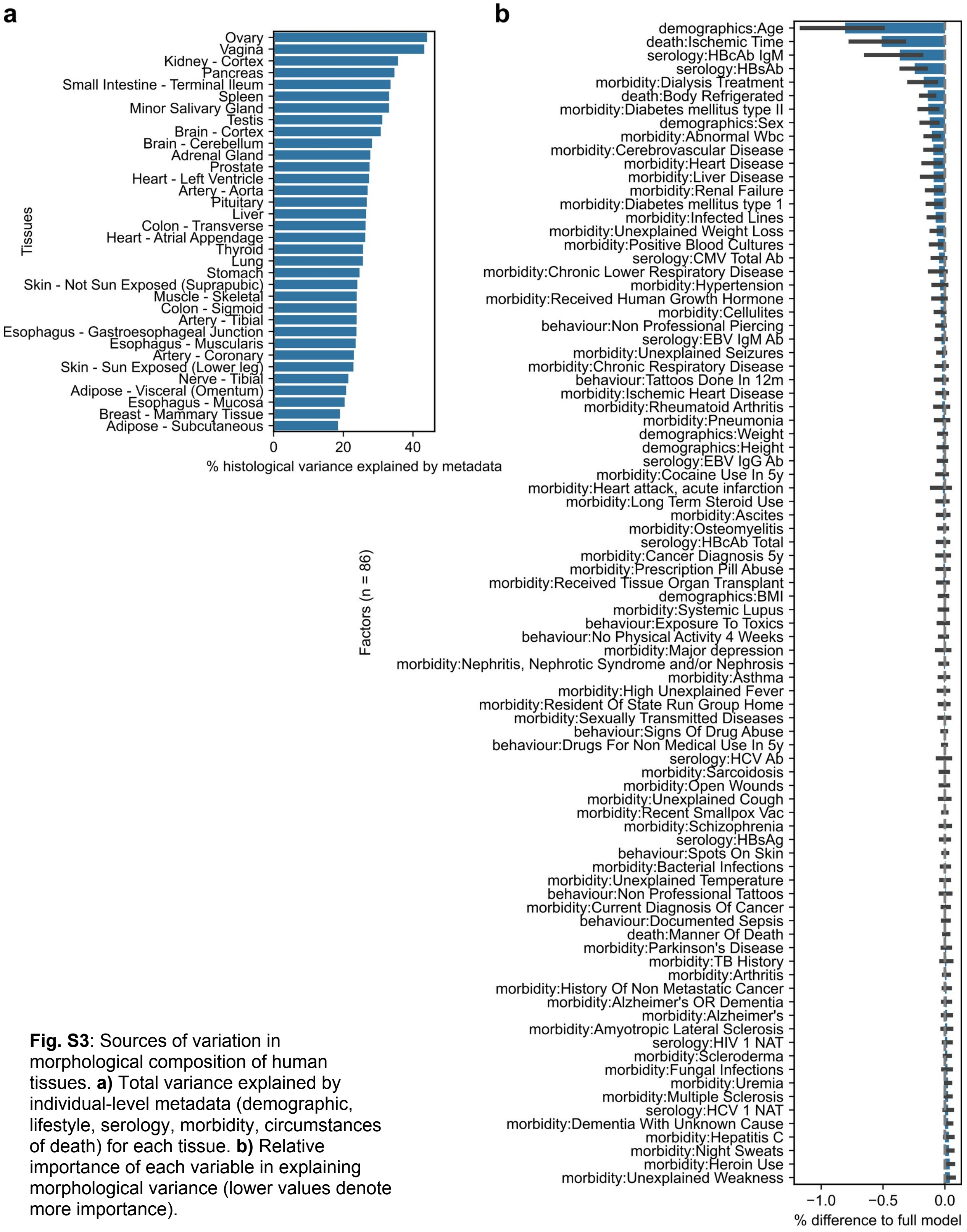


**Fig. S3**: Sources of variation in morphological composition of human tissues. **a)** Total variance explained by individual-level metadata (demographic, lifestyle, serology, morbidity, circumstances of death) for each tissue. **b)** Relative importance of each variable in explaining morphological variance (lower values denote more importance).

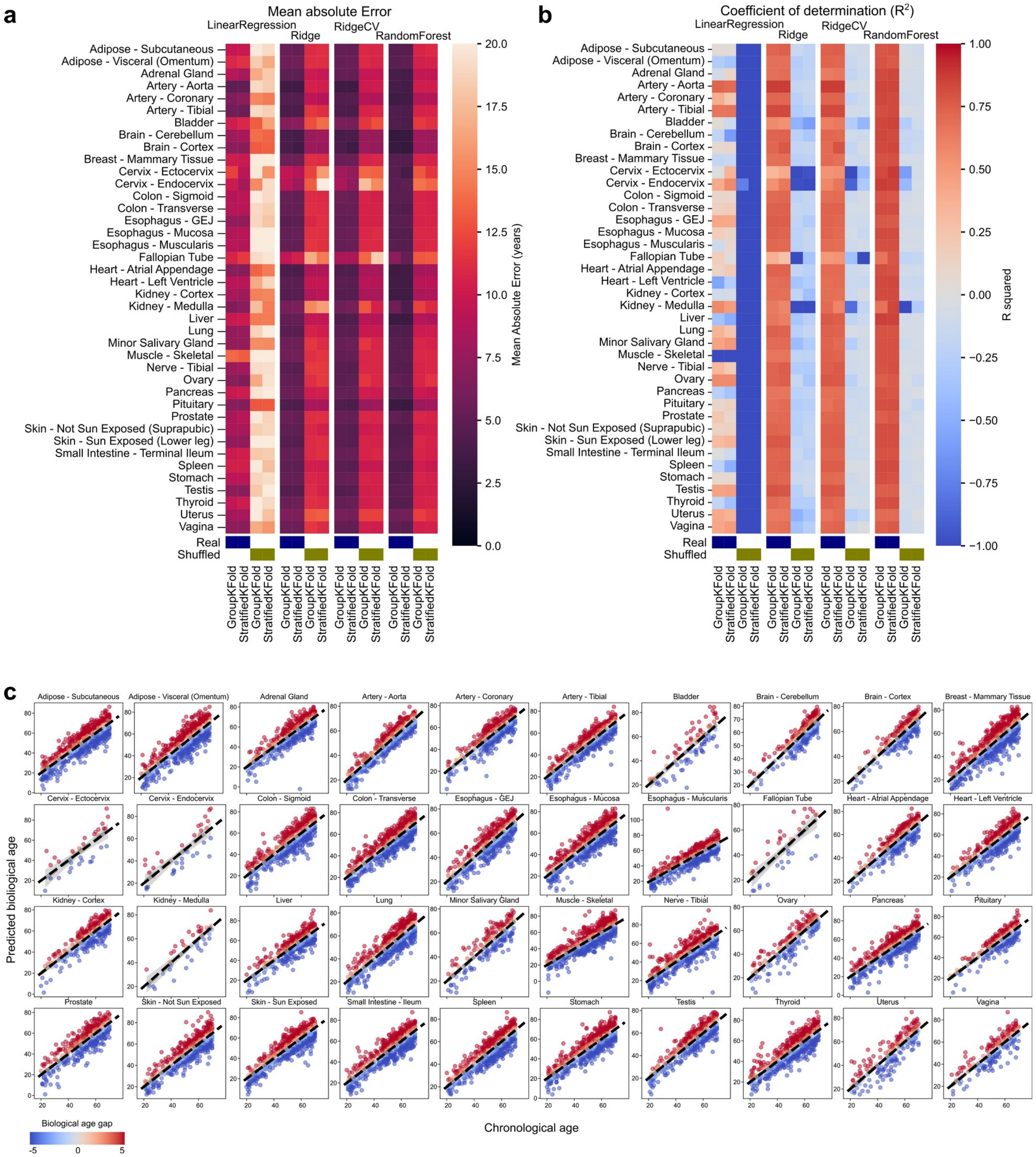


**Fig. S4**: Performance of tissue clocks. **a-b)** Assessed with the mean absolute error (a) and coefficient of determination (b) metrics for each tissue using different predictors and cross validation strategies. **c)** Visualization of the predicted biological age (y-axis) contrasted with chronological age (x-axis) of each individual in each tissue.

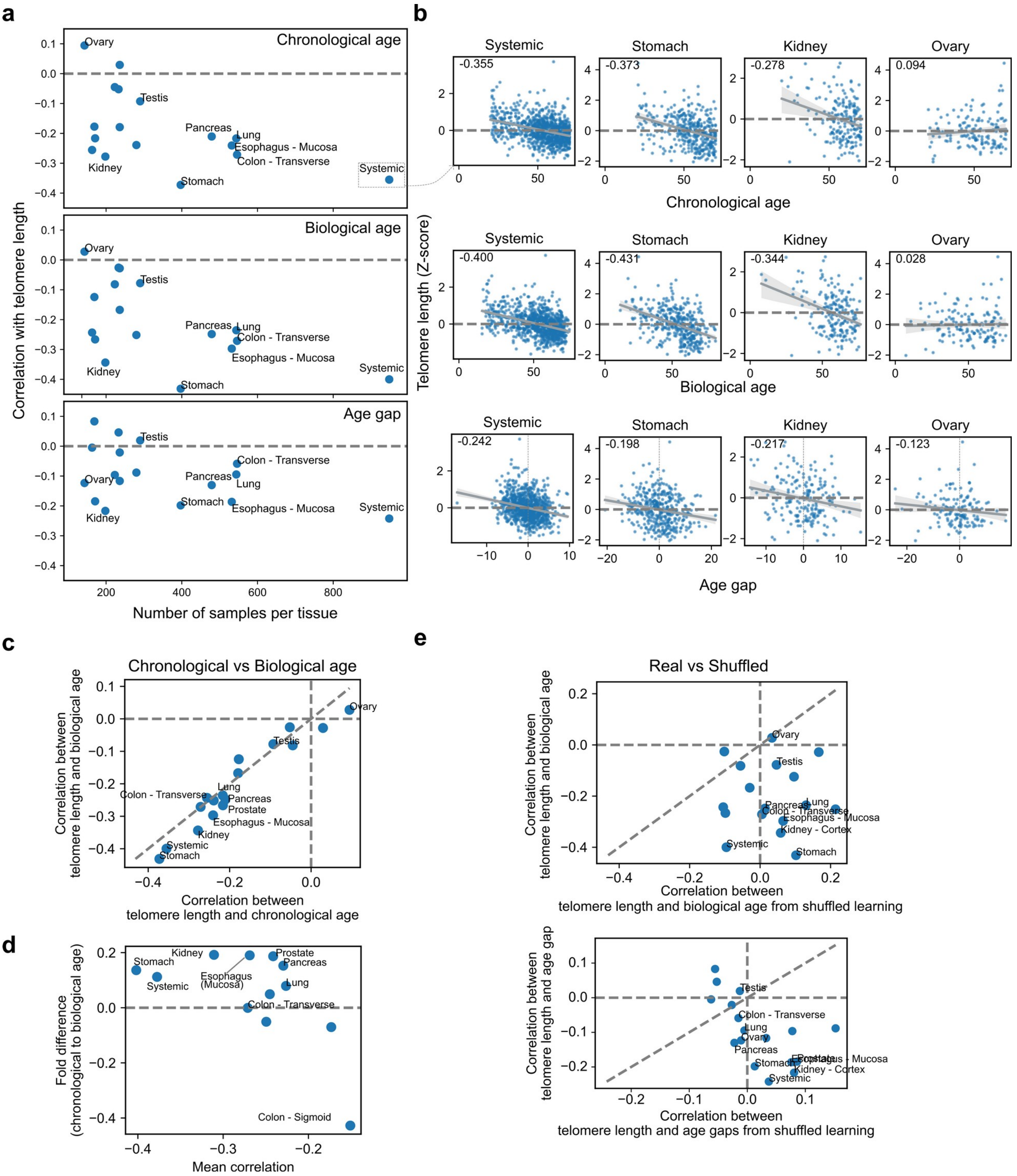


**Fig. S5**: Relationship between inferred age gaps and telomere lengths measured in the matched samples. **a-b)** Relationship between chronological age (top), biological age (middle) and age gaps (bottom) and the length of telomeres per tissue, dependent on the number of samples available for each tissue (a). Specific examples are also shown (b). **c)** Comparison of chronological age and biological age in the association with telomere lengths per tissue. **d)** As in c) but highlighting the fold change in Pearson correlation with telomere lengths between biological and chronological age in particular for tissues with high sample size. **e)** Comparison of predictors trained on real vs shuffled target labels (age) in the association with telomere lengths.

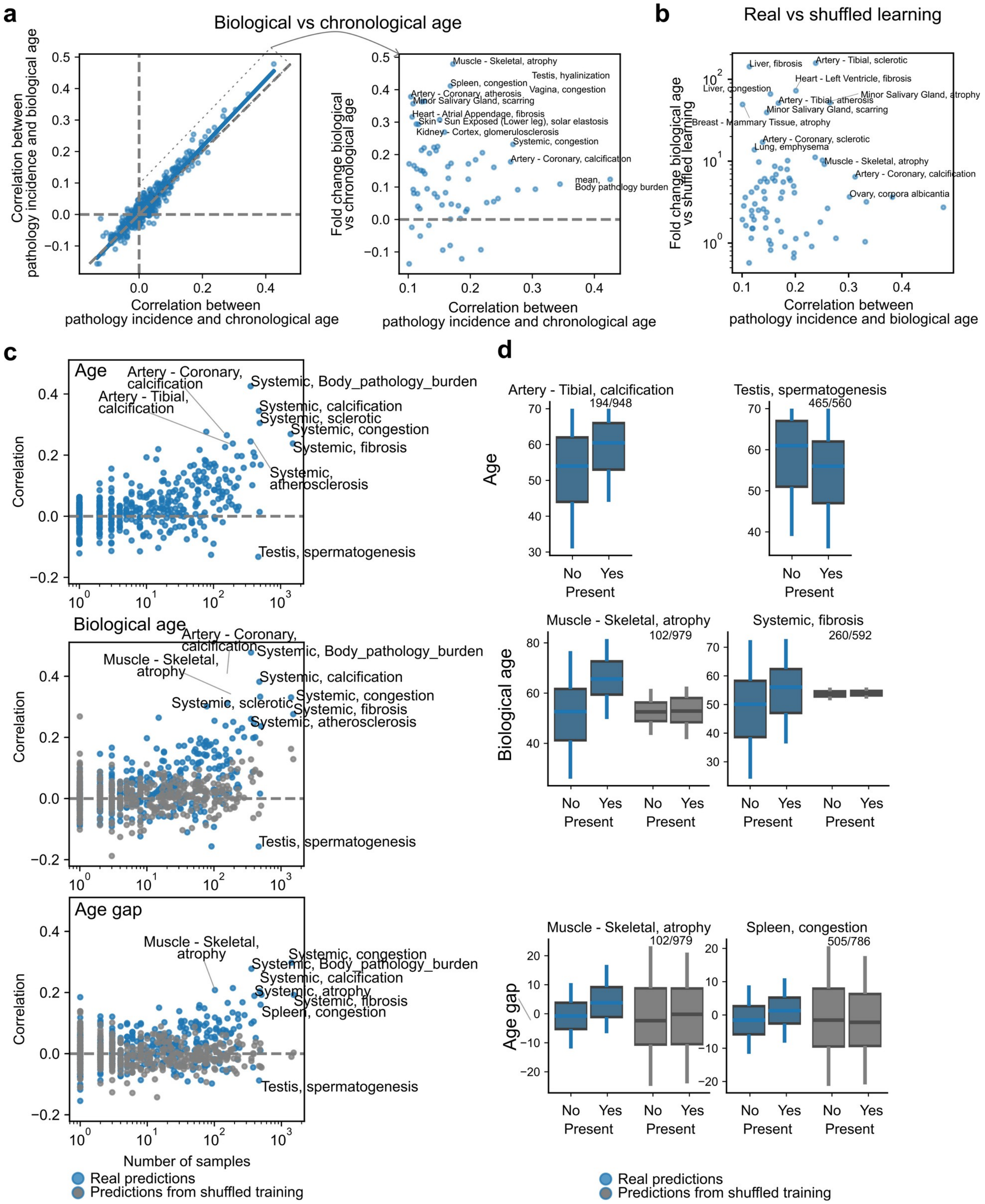


**Fig. S6**: Relationship between the predicted biological age and incidence of tissue pathology. **a)** Comparison of the association of biological age vs chronological age with tissue-specific pathologies. The dashed line represents x = y, and the box is highlighted in the right as a fold increase in the association with pathology for biological vs chronological age. **b)** Comparison of biological age associations with pathology in real vs shuffled learned predictions. **c)** Pearson correlation between the incidence of tissue specific pathology and chronological age (top), biological age (middle) and age gaps (bottom). Each point is a pair of pathology type in a tissue. Blue points are estimates from predictors using real data, and grey points from predictors trained on shuffled target label (age) as control. **d)** Examples of pathologies affecting specific tissues with differential associations with age.

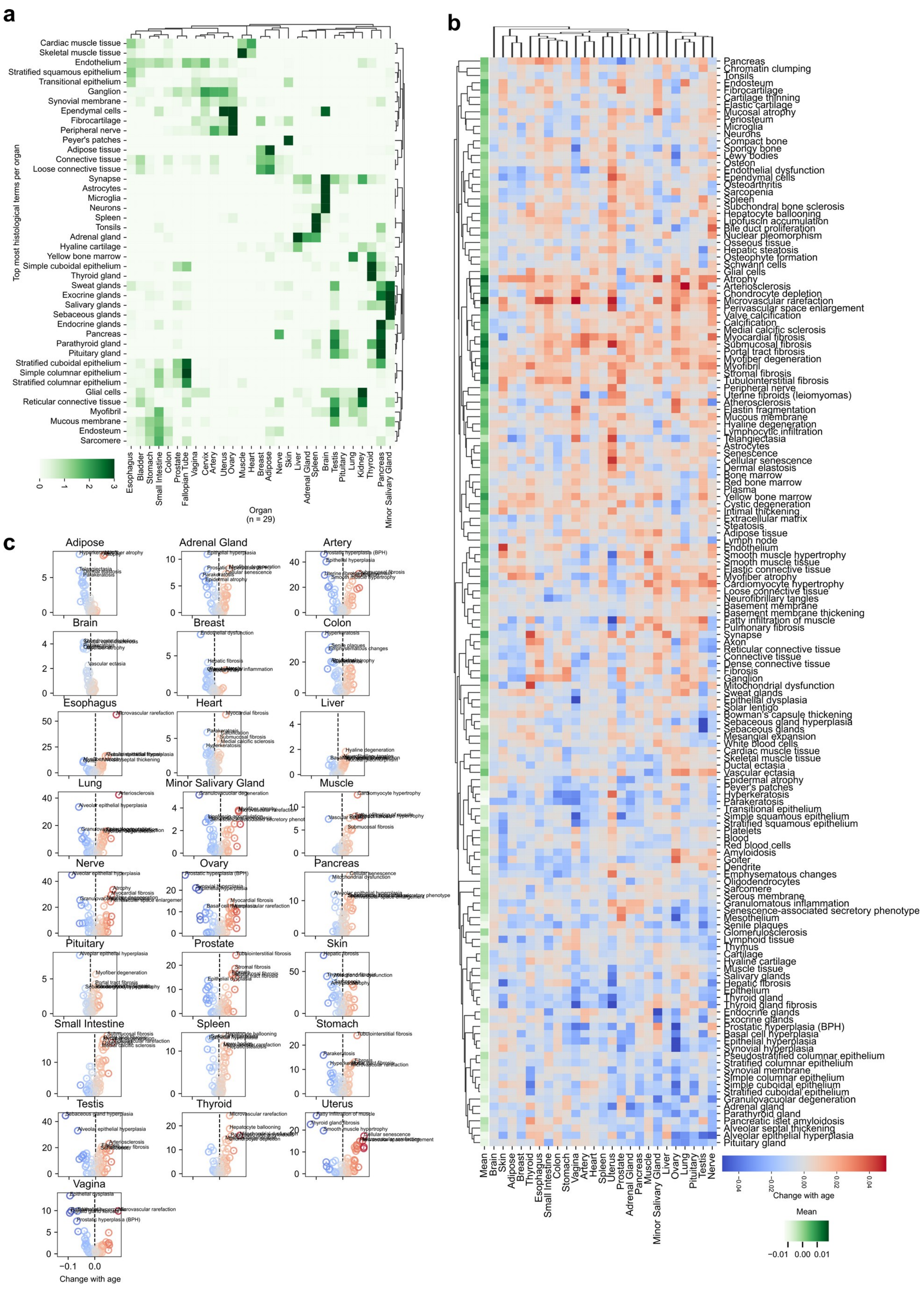

**Fig. S7**: Characterization of histological features associated with age gaps using multi-modal text-image models. **a)** Heatmap with the terms with highest value for each organ. **b)** Heatmap of changes in text terms with biological age estimates. **c)** Same as B) but as a volcano plot where the y-axis is the FDR-adjusted -log10(p-value).

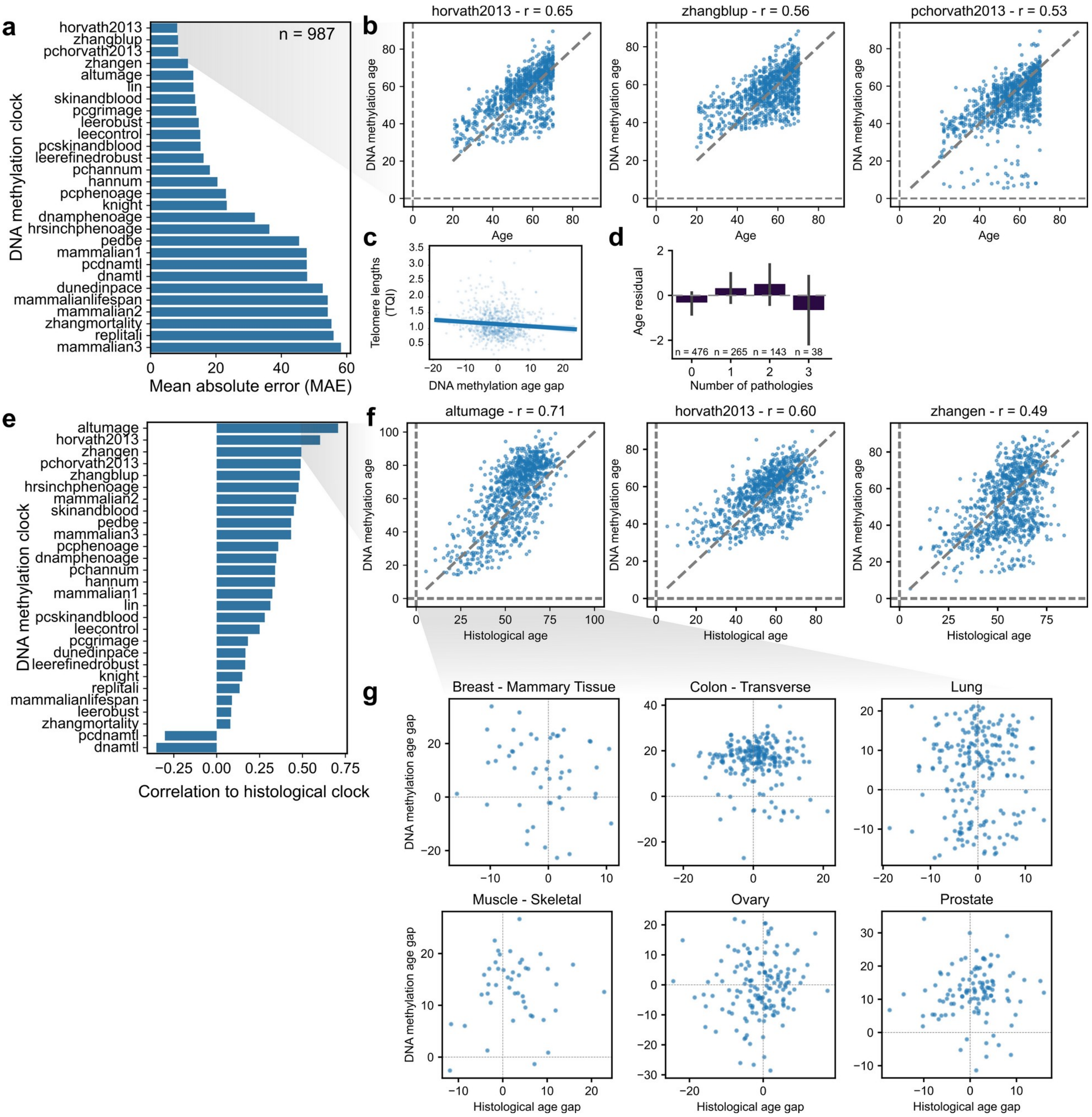


**Fig. S8**: Comparison of morphological tissue clocks to DNA methylation clocks. **a)** Mean absolute error for a variety of established DNA methylation clocks applied to the GTEx cohort (987 samples). **b)** Illustration of the 3 best performing DNA methylation clocks. **c)** Relationship between predicted age gaps using the Horvath2013 DNA methylation clock and telomere lengths of the same tissue samples. **d)** Enrichment of pathological terms dependent on estimated DNA methylation age gap. **e)** Pearson correlation values of DNA methylation and histological clocks. **f)** Illustration of the predictions of 3 most well correlated DNA methylation clocks and the predicted biological age from histological clocks. **g)** Relationship between predicted age gaps in the AltumAge DNA methylation clock and the age gap from histology.

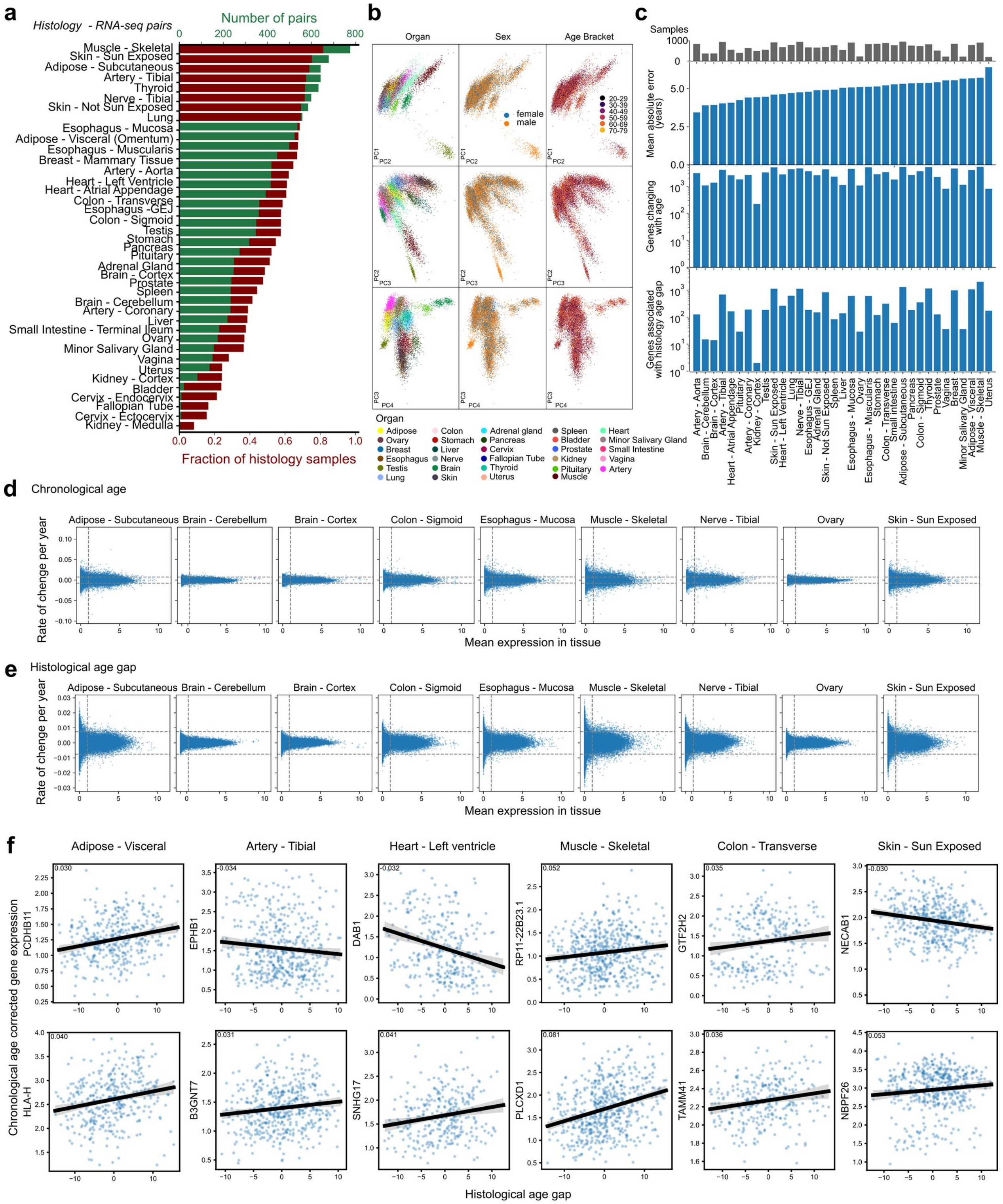

**Fig. S9**: Molecular basis of histological age associated changes in 40 human tissues. **a)** Number and proportion of histological samples with matched RNA-seq data. **b)** Principal Component Analysis of RNA-seq data. **c)** Comparison across tissues in WSI-RNA-seq sample pairs, tissue-clock performance, and number of genes associated with chronological age and histological age gaps. **d-e)** Relationship between tissue-specific mean expression and age-associated rates of expression changes (MA plot) for chronological age (c) and histological age gap (d). **f)** Examples of genes changing with histological age gap alone – to illustrate this effect chronological age was regressed out from the data.

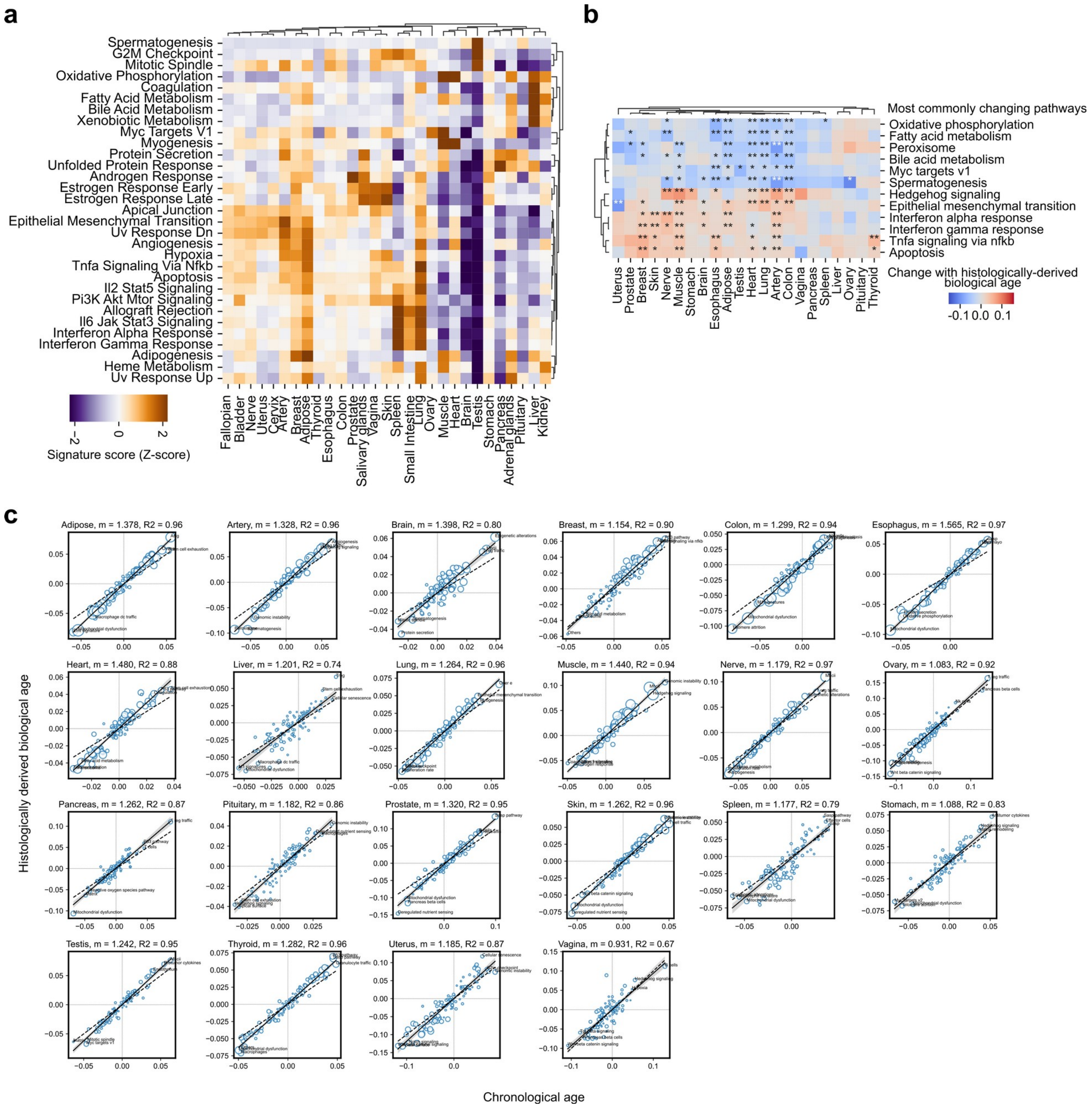


**Fig. S10**: Comparison of transcriptional changes with chronological and biological age derived from histology. **a)** Heatmap with top MSigDB Hallmark pathways for each organ. **b)** Heatmap with top and bottom 6 most consistently up- or down-regulated pathways across organs. **c)** Relationship between changes associated with predicted age gaps from histological images (y-axis) and chronological age (x-axis).

a

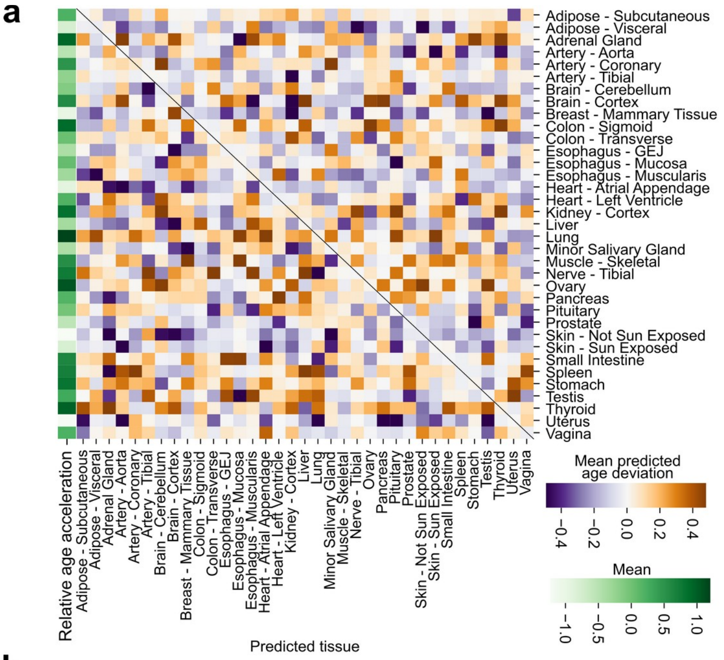


**Fig. S11**: Tissue clock cross-prediction between tissues. **a)** Values indicate the mean age gap predicted from the tissues in the rows in the tissues in the columns. **b-c)** Prediction of biological age (b) and its associated age gap (c) from multiple tissues (y-axis) in each tissue (panels). The black line is a rolling mean with a decade window. The orange line a 3rd order spline regression. The blue line is a lowess fit.

b

c

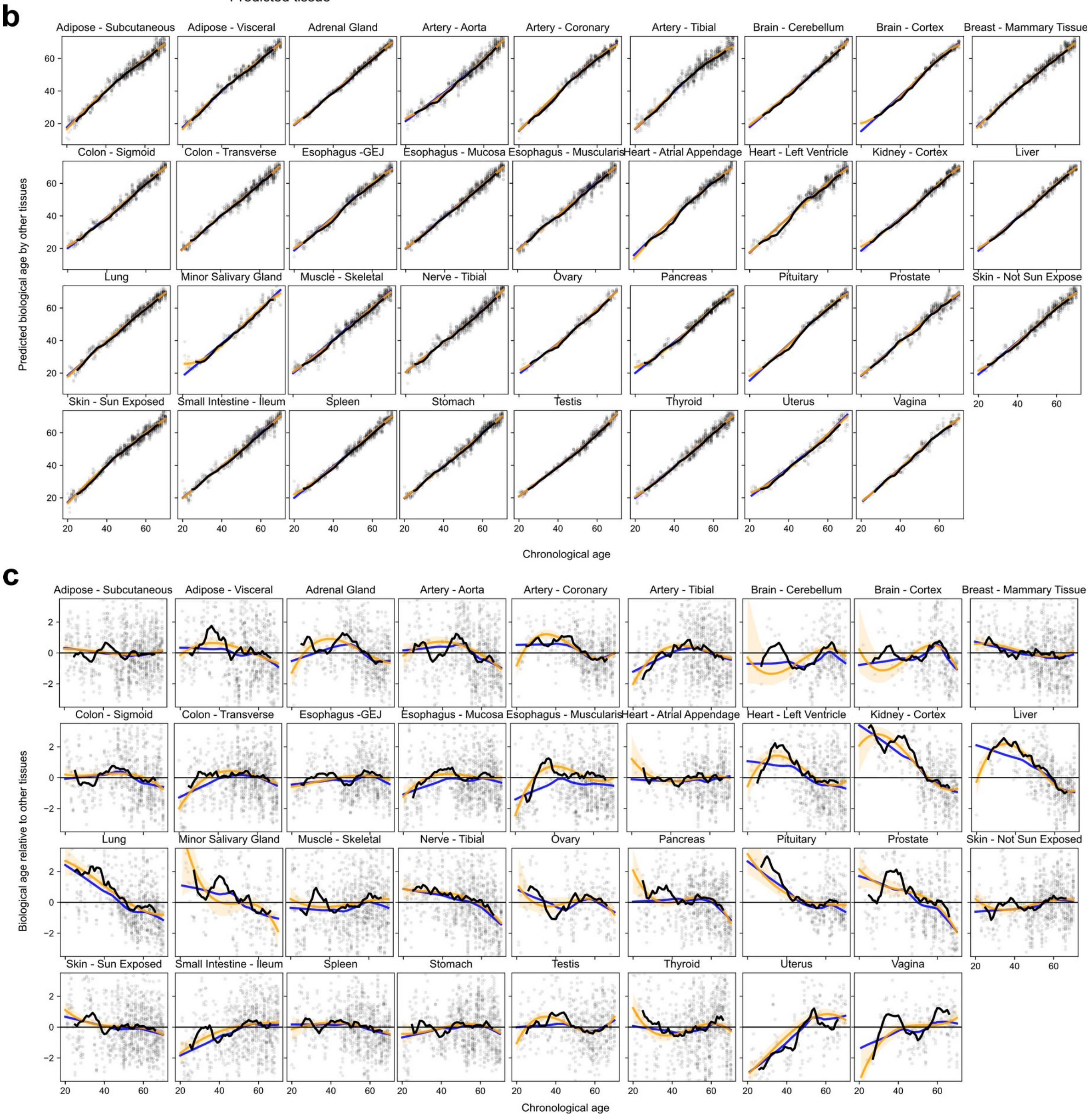

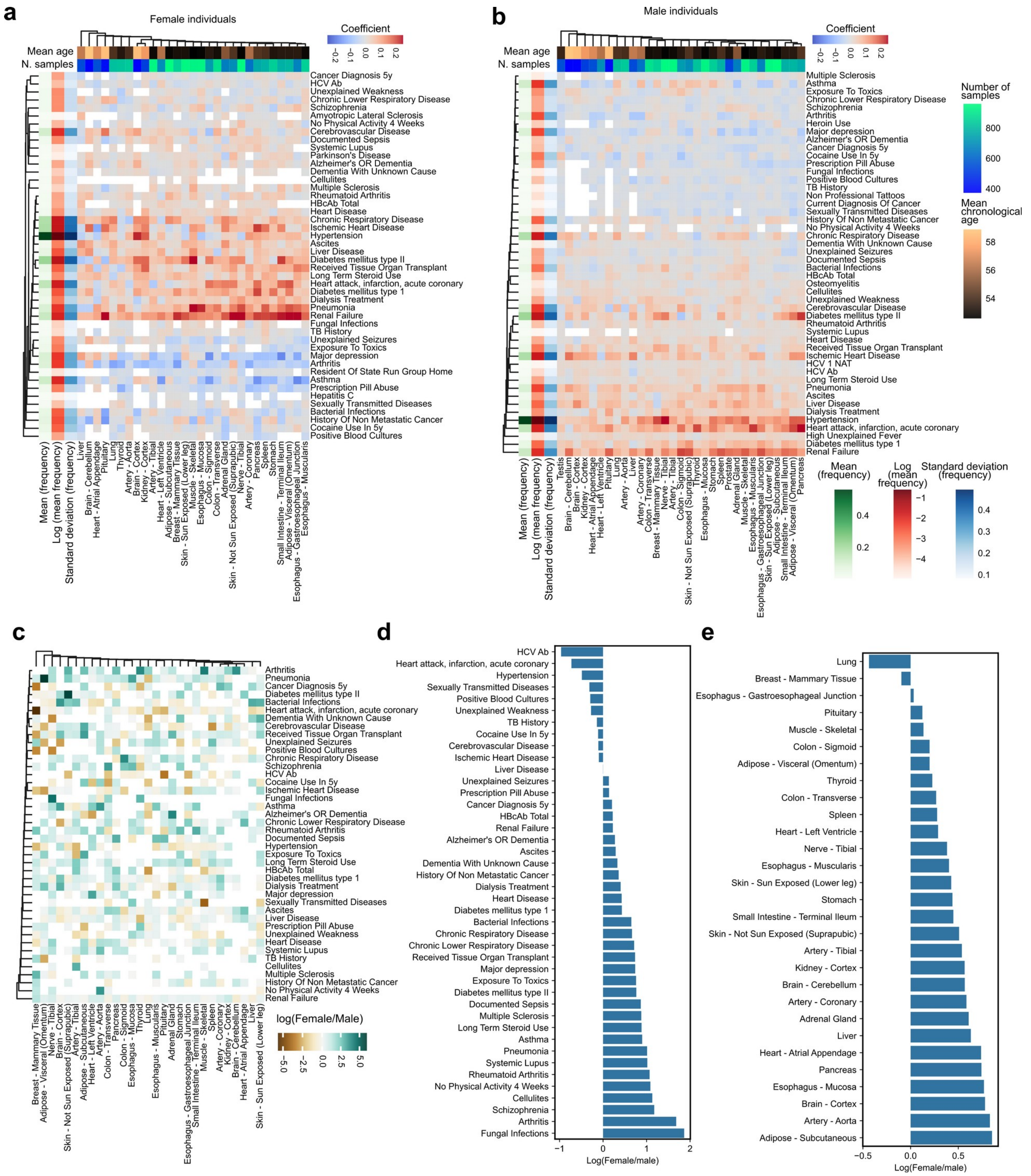


**Fig. S12**: Association of tissue-specific age gaps with factors. **a-b)** Coefficients of association between tissue-specific age gaps and known factors from female a) or male b) individuals. **c)** Log fold enrichment of effects between female and male individuals. **d-e)** Log fold enrichment between female and male individuals in the overall association with known factors d) and tissues e).

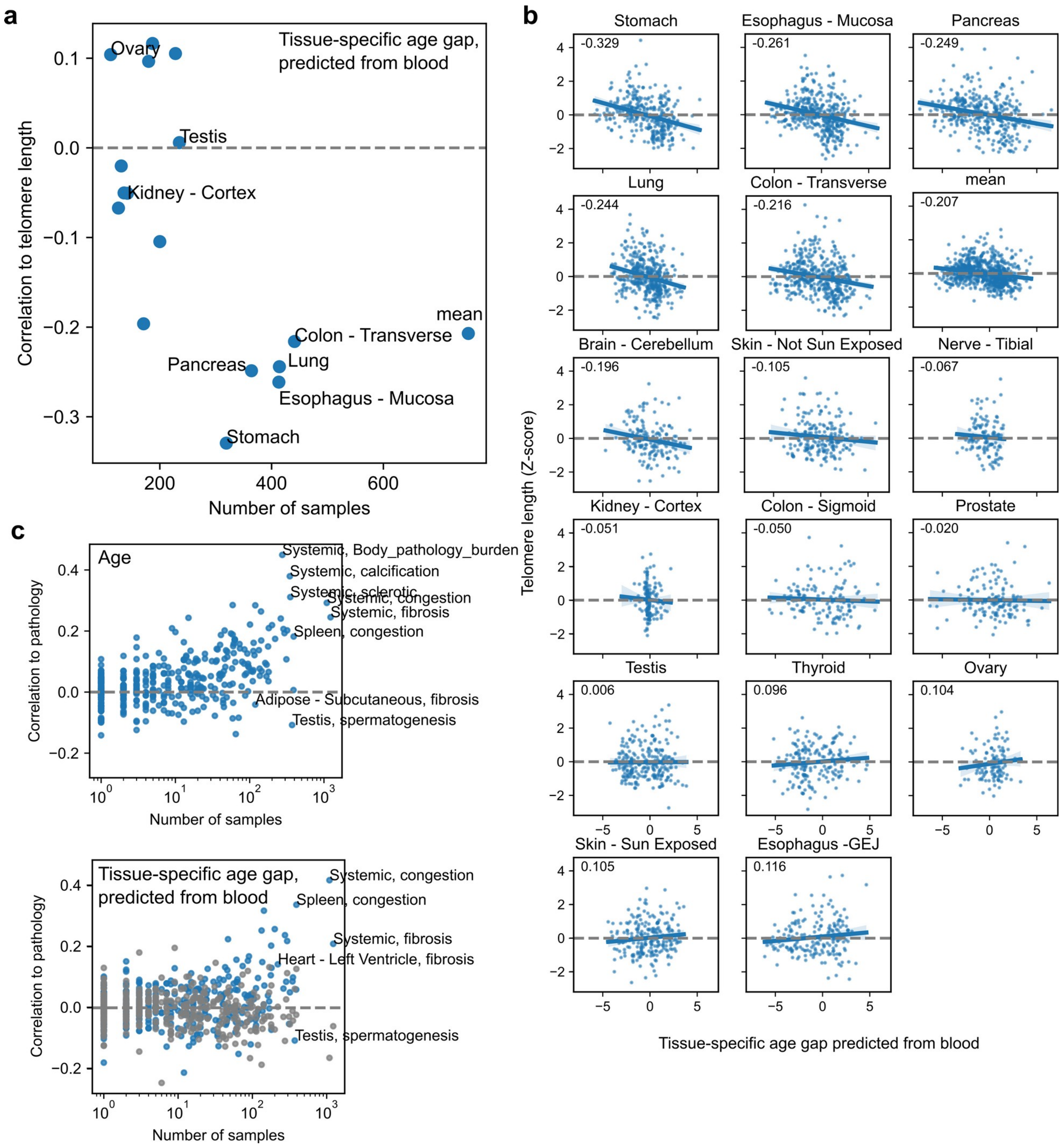


**Fig. S13**: Association of tissue-specfic age gaps predicted from blood with telomere lengths and pathology incidence. **a)** Correlation between blood-based predicted tissue-specific histological age gaps and telomere lengths for each tissue. **b)** As in a) but illustrating the relationship for each sample in each tissue. The Pearson correlation value is indicated in the upper left corner of each plot. **c)** Correlation between chronological age (top) or blood-based tissue-specific histological age gaps (bottom) with incidence of tissue specific pathology. Each point is a tissue-pathology pair.

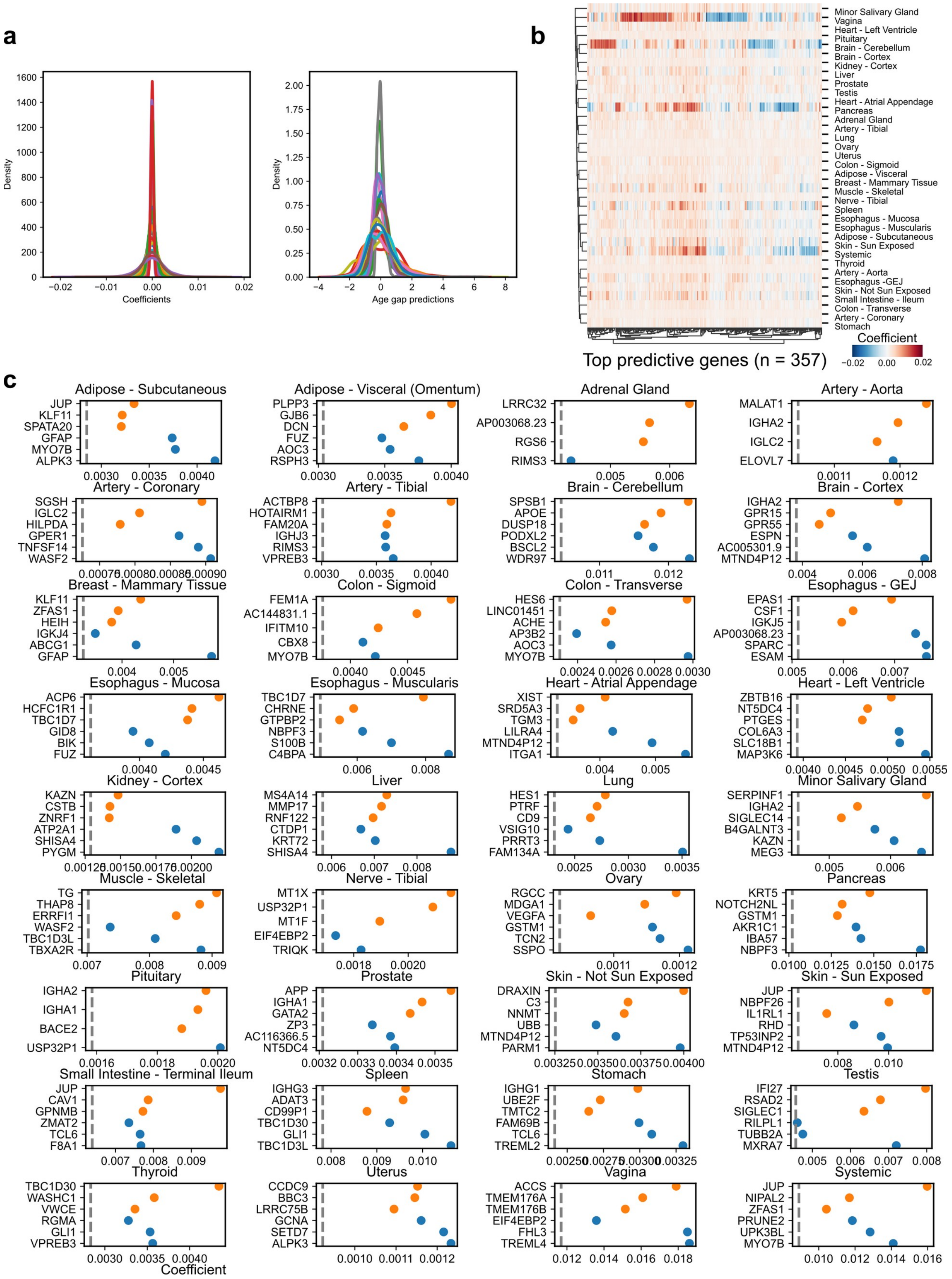


**Fig. S14**: Interpretation of the prediction of tissue-specific age gaps in blood gene expression. **a)** Histograms of estimated coefficients (left) or model residuals (right) for each organ. **b)** Clustered heatmap with set of predictive genes across tissues. **c)** Top 3 predictive genes for each tissue. Orange points are genes positively associated with age gaps, whereas blue points are negatively associated – note that the coefficient sign was inverted for negative associations to simplify visualization. The dashed gray line is the 99.9[th] percentile of absolute coefficients for each tissue clock.

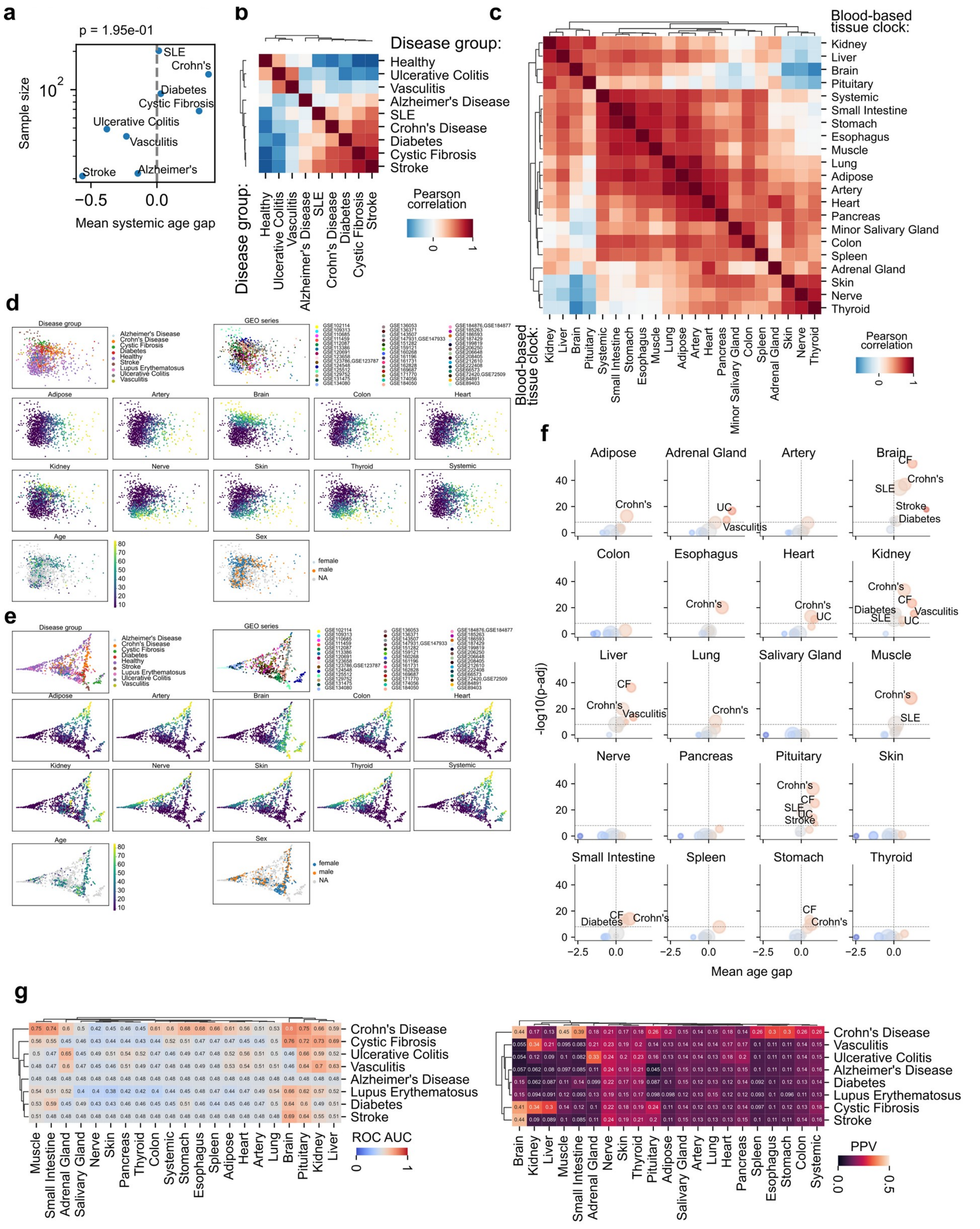

**Fig. S15**: Validation of blood-based tissue-specific clocks in independent cohorts. **a)** No association between systemic age gap and sample size of each disease group. **b-c)** Pairwise correlation of diseases (b) or organs (c) across predicted age gaps. **d-e)** PCA (d) and diffusion maps (e) based on predicted age gaps for all samples. Each plot is colored by disease, study, age, sex, as well as the tissue-specific age gap. **f)** Volcano plots for each organ showing the statistical association of age gaps for each disease compared with healthy samples (t-test, Benjamini-Hochberg FDR correction). **g)** Thresholded values of age gaps were used to asses the area under the receiver operator curve (ROC AUC) and positive predictive value (PPV) of the blood-based tissue-specific predictors.
